# Reciprocal regulation of autophagy and exosome pathway is mediated by GABARAPL2 and Alix to facilitate cellular homeostasis

**DOI:** 10.1101/2024.11.02.621656

**Authors:** Naveen Soni, Megha Chaudhary, Bhawana Bissa

## Abstract

The continuous reliance of cancer cells to acquire energy and communicate their nutrient needs makes them resilient and vulnerable. It provides an opportunity to stifle cancer cells by restricting their energy generation and communication ability. Autophagy and exosome biogenesis are two such pathways that are essential in maintaining the robust growth and survival of cancer cells. In this study we observed that inhibition of one pathway altered the expression of genes in other pathway. Exosome biogenesis, when blocked, led to an increase in breast cancer cell proliferation, while inhibition of autophagy did not significantly affect cancer cell proliferation. Therefore, the two pathways, when independently inhibited, did not present any significant effect on restricting cancer cell growth. However, we observed a substantial reduction in cancer cell proliferation upon combined inhibition of two pathways. To evaluate the reciprocal regulation of two pathways, we blocked the autophagy pathway and observed increase in the secretion of exosomes from MDA-MB-231 cells, along with decreased expression of Alix and CD63. On contrary, inhibition of exosome biogenesis led to an increase in the expression of ATG5 and ATG16L1, which caused a significant decrease in expression of GABARAPL2. Interestingly, the knockdown of GABARAPL2 abrogated the decrease in Alix expression upon autophagy inhibition, thus highlighting the essential role of GABARAPL2 in Alix secretion. Thus, our study highlights for the first time the synergistic effects of autophagy and exosome pathway inhibition in restricting cancer cell growth as well as the involvement of GABARAPL2 in the regulation of exosome secretion via modulating Alix expression.

## Introduction

Extracellular vesicles (EVs) are micro/nano-sized monolayered secretory vesicles that are secreted by almost all types of cells and are a conserved mechanism throughout the evolution[1], [2], [3]. Initially, in 1983, they were discovered as secretory vesicles from immature RBCs in sheep but later termed “exosomes” in 1987[4], [5]. Exosomes originate from the endosome, in which the inward budding of the endosome membrane causes the formation of intraluminal vesicles (ILVs), which later mature into multivesicular bodies (MVBs)[6], [7]. The membrane proteins and incorporated cytosolic proteins in the exosomes are generally tetraspanins (CD9, CD63, CD81), Alix, TSG101, HSPs, Actin, and Flotillin[8], [9]. Till now, two processes have been identified for exosome synthesis, i.e., an endosomal sorting complex required for transport (ESCRT) dependent and ESCRT independent. The discovery of ESCRT proteins in cargo loading and membrane modeling was a breakthrough mechanism that uncovered the mystery behind MVB synthesis[10]. ESCRT machinery proteins ESCRT-0, ESCRT-I, ESCRT-II, ESCRT-III, and VPS4 are present on the cytosolic surface of MVB membrane and participate cooperatively to promote vesicles budding, cargo sorting, and exosome synthesis[11], [12], [13], [14]. This machinery works in a stepwise cascade in which ESCRT-0 and ESCRT-I first initiate the clustering of ubiquitylated cargoes onto the microdomains present on the MVB membrane. This recruits ESCRT-II and ESCRT-III, which follow the inward budding of membrane and fission of microdomains, resulting in a flask-shaped bud inside the MVB lumen. Alix, another critical component of exosome machinery, assists budding and ILV formation by binding with the ESCRT-III subunit[15]. An ATPase-VPS4 finally buds off the vesicle[16]. ESCRT-Independent exosome synthesis was discovered when CD63-loaded exosomes were collected from the ESCRT-depleted cells[17]. The ESCRT-independent mechanism can be elaborated as ceramide-dependent and tetraspanin-dependent. In the former, sphingomyelin is hydrolyzed by neutral sphingomyelinases (nSmases) to generate ceramide that may generate membrane subdomains and make a membrane curvature inside the MVBs[18], [19]. While tetraspanin-dependent, CD63 is mainly thought to regulate the endosomal sorting mechanism during melanogenesis in melanocytes[20], [21]. Additionally, proteins such as CD9, CD63, CD81, and CD82 also assist cargo sorting to the exosomes. Although both mechanisms are present in the cells, their contributions to generating exosomes may depend on the cell and cargo types.

Autophagy is a highly regulated process that is present at the basal level in most of the cells, but sometimes, it can be used as a survival strategy by cancer cells. Autophagy has a dual role in tumor progression and suppression[22], [23]. It has been classified into three types: Macroautophagy, Chaperone-mediated autophagy (CMA), and Microautophagy. The autophagy process includes the initiation of phagophore formation, nucleation, and elongation of the autophagosome, as well as its fusion and degradation in the lysosome[24]. Briefly, under stress conditions, the mammalian target of rapamycin (mTOR) gets inactivated by AMPK and TP53, which leads to the activation of the ULK complex and ATG13, i.e., initiation of autophagy[25]. In the nucleation step, another protein beclin-1 interacts and conjugates with VPS34 and ATG14L at the phagophore membrane to promote the recruitment of protein turnover and damaged organelles[26], [27]. In the elongation and maturation of the autophagosome membrane, ATG8 plays a crucial role in intracellular trafficking. ATG8 can be subdivided into two protein subfamilies, i.e., microtubule-associated protein light chain-3 (MAP1LC3A-C/LC3A-C) subfamily and γ-aminobutyric acid receptor-associated protein (GABARAP, GABARAPL1, GABARAPL2/GATE-16) subfamily[28], [29]. While the MAP1LC3 system regulates initial phagophore membrane elongation, GABARAB proteins are thought to be involved in the final sealing of autophagosome membrane[30]. These subfamilies are cleaved by two different protein systems. The first system involves ATG4, ATG7, and ATG3 and the second system is ATG12, ATG7, ATG16L1, and ATG5. These complexes interact and produce lipidative forms of LC3 and GABARAP subfamilies. Later, the autophagosome fuses with the lysosome to form autolysosome, and cargo is degraded.

GW4869 is a non-competitive inhibitor for the nSmase enzyme, which was first used in HEK293 cells to inhibit exosome release[31], [32]. During ESCRT-independent exosome synthesis, the cone-shaped structure of ceramide plays a crucial role in the formation of lipid raft domains, which triggers the construction of a negative curvature onto the MVB membrane[18]. Therefore, inhibition of nSmase by GW4869 reduces the release of exosome by ESCRT-independent pathway, thus lowering the overall secretion.

Chloroquine (CQ or CHQ) was a well-known antimalarial drug, but after developing resistance against CQ in malarial parasites, its use has been diminished[33]. Previous studies on the use of CQ in cancers have determined that CQ has the potential to increase radiotherapy and chemotherapy sensitivity in a broad range of cancers[34]. CQ diffuses in the cells in unprotonated form; since it is a weak base, it gets accumulated and becomes protonated in the lysosomes[35]. Consequently, it raises intralysosomal pH and disturbs autophagosome-lysosome fusion, thus inhibiting autophagy. Inhibition of autophagy results in a stock of damaged organelles in the cytoplasm and protein turnover in the ER, causing ER stress. Wortmannin was identified as the first phosphatidylinositol-4,5-biphosphate 3-kinase (PI3K). Later, it was discovered that it interacts with additional PIEK accessory proteins, indicating potential adverse effects if produced as a drug. In this study, we identified the change in size and intensity of exosomes derived from CQ-treated MDA-MB-231 cells. We also investigated the crosstalk between exosome and autophagy pathways using CQ and GW4869 in breast cancer cell line MDA-MB-231, and HEK293T fibroblast cells. Furthermore, for the first time, we have identified the link between cross-regulation of GABARAPL2 and Alix in mediating the homeostasis between Autophagy and Exosome pathways.

## Results

### 1. Autophagy inhibition enhances exosome release in breast cancer cells

To determine whether inhibiting autophagosome-lysosome fusion can alter the exosome pathway, 20µM CQ was added and after 24 hrs cell culture media (CCM) of MDA-MB-231 cells was collected. Exosomes were isolated from an equal volume of CCM in the control and CQ-treated MDA-MB-231 cells and MCF-7 cells. Using differential light scattering (DLS), an increase in exosome size was observed after CQ treatment in both MDA-MB-231 (Supp. Fig. 1A) and MCF-7 cells (Supp. Fig. 1B). As per previous studies, a change in the physical properties of exosomes, such as increased exosome diameter (Fig. 1A) and intensity (Fig. 1B) from CQ-treated MDA-MB-231 cells were observed by nanoparticle tracking analysis (NTA) [36]. The data presented the mean value for the exosomes of control MDA-MB-231 cells to be 149nM ±58.4 and a concentration of 4.57e+007±1.32e+007 particles/mL (6.2±1.3 particles per frame and 6.7±1.4 centres per frame) (Fig. 1C). Meanwhile, exosomes from CQ treated MDA-MB-231 cells presented a mean value of 169.5nm±95.6 and a concentration of 5.62e+007±4.78e+006 particles/mL (10.0±1.7 particles/frame and 10.3±1.6 centres per frame) (Fig. 1C). To further characterize exosomes, the same set of exosome samples were imaged under Transmission electron microscopy (TEM) with a magnification of 1,00,000x at 100nm scale. A homogeneous population of rounded vesicles with 25-150nm was observed, indicating that the exosomes were intact and not ruptured during the isolation (Fig. 1D). As per the previous studies, we further performed SDS-PAGE (Supp. Fig. 1C) and immunoblot assay to detect exosome-specific biomarkers in exosome lysates[13], [37], [38]. The exosomes showed the presence of TSG101, β-actin, and CD63 (Fig. 1E) with a relatively high expression of β-actin and TSG101 in exosomes isolated from CQ-treated MDA-MB-231 cells (Supp. Fig. 1D).

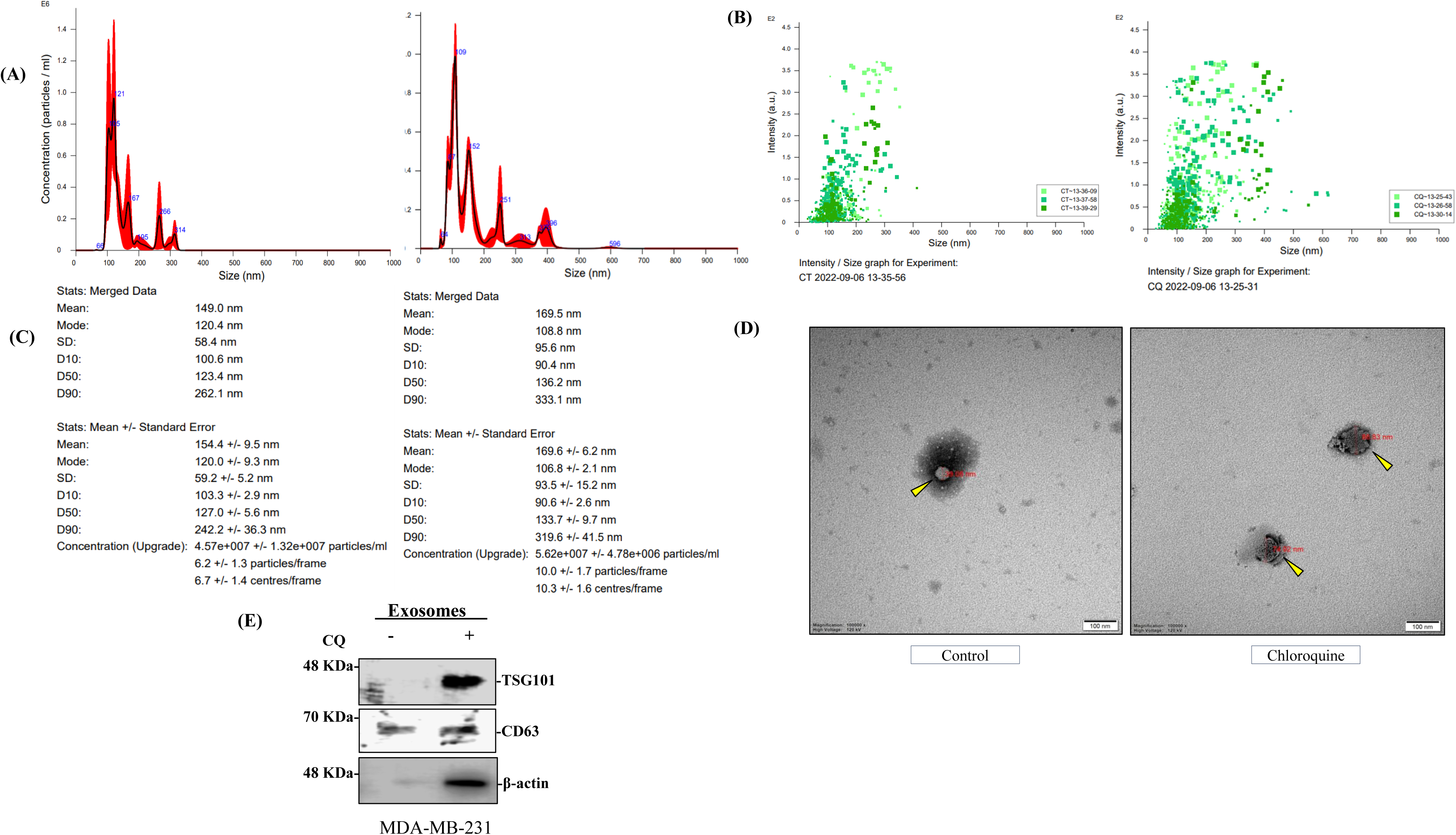

### 2. Autophagy inhibition decreases the expression of exosome-specific markers in cells while increasing the release in exosomes

To evaluate the effect of autophagy inhibition on exosome pathway we treated MDA-MB-231 cells with 20µM CQ for 24 hrs and observed that both CD63 and Alix mRNA expression was downregulated (Fig 2A). We further checked the effect on protein levels, and similar to previous studies [39], we found exosome biomarkers Alix, as well as flotillin, to be decreased in the CQ-treated MDA-MB-231 cell lysates but consequently increased in the exosomes from CQ-treated MDA-MB-231 cells (Fig. 2B). A most recent study has shown that tumor exosomes influence nearby and distant cells by remodeling microenvironment and drug resistance[40], [41]. We incubated MDA-MB-231 cells with the exosomes derived from control and CQ-treated MDA-MB-231 cells to check how cancer exosomes affect the survival of tumor cells. This result showed a significant cell death in cells treated with CQ exosomes compared with control conditions (Supp. Fig. 2A, 2B). In addition, we also detected a decrease in Alix in the CQ treated MDA-MB-231 and HEK293T cells but no such changes were observed in CD63 and TSG101 (Fig. 2C, 2D). The decrease in Alix was significant in both MDA-MB-231 and HEK293T cells, so we treated HEK293T cells with CQ in a time-dependent manner and observed a gradual decrease in Alix expression (Supp Fig. 2D). Previous studies have shown that Rab27a and Rab27b promote exosome secretion in different cancers[42], [43]. We observed a higher expression of Rab11 and Rab27a in CQ-treated HEK293T cells (Fig. 2E), which denotes an up-regulation of exosome secretion upon blocking autophagy flux. The illustration (Supp. Fig. 2E) depicts the role of different Rab proteins in the co-regulation of both autophagy and exosome pathways.

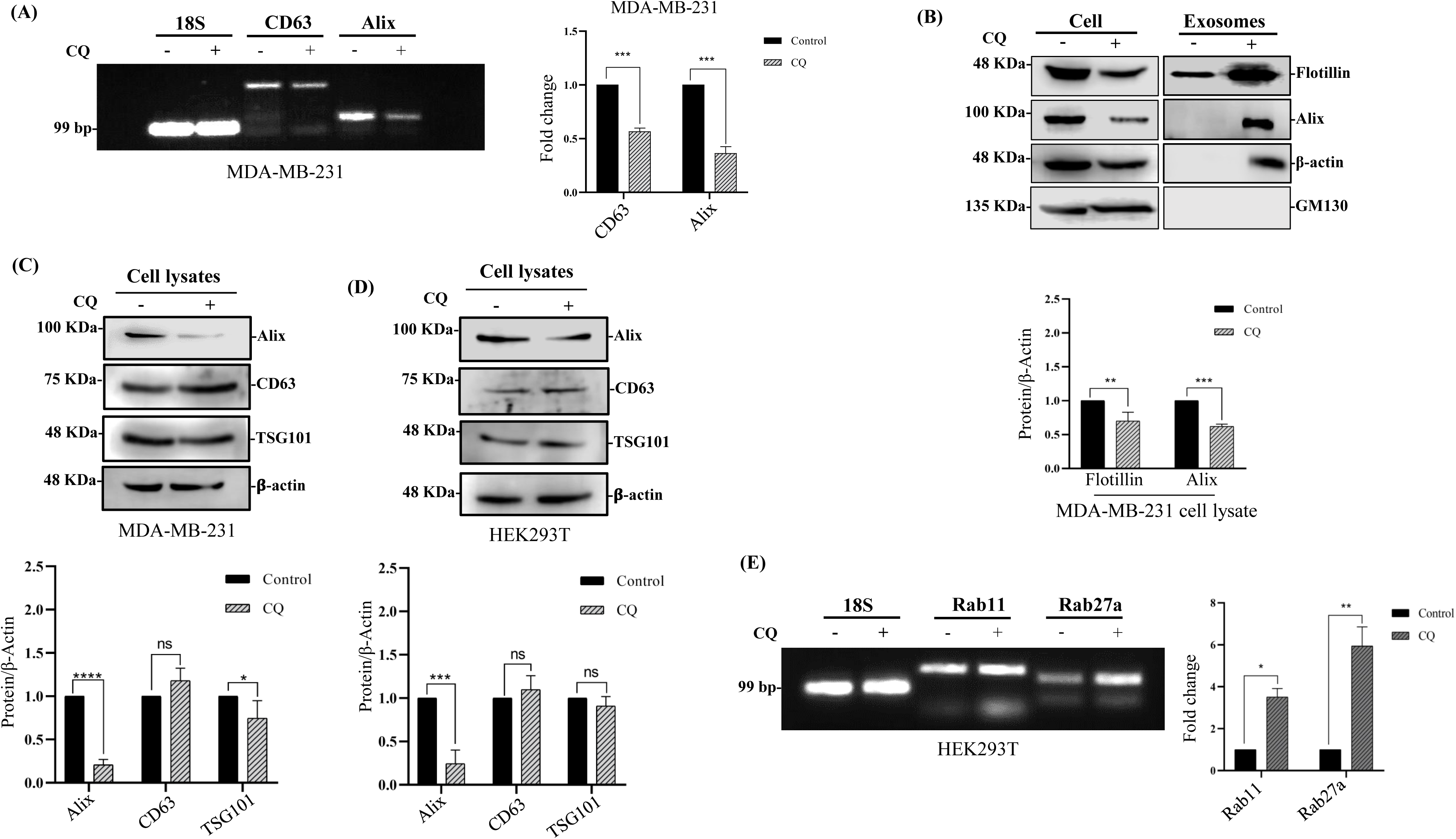

### 3. Exosome pathway inhibition alters the expression of autophagy-related genes

A recent study has shown the role of ATG5 in MVBs acidification[44] and another study has demonstrated that silencing of ATG5 enhances exosome release[45], so we wanted to analyze the expression of autophagy genes after inhibiting exosome biogenesis by using an inhibitor of nSmase2, GW4869 treatment in MDA-MB-231 cells. We observed an increase in ATG5 (>3 fold) and ATG16L1 mRNA expression (>2 fold) after treatment with 10µM GW4869 in MDA-MB-231 cells for 24 hrs (Fig. 3A, 3B). This increase in ATG5 mRNA expression was observed both in MDA-MB-231 and MCF-7 cells (Supp. Fig. 3A). Protein expression of ATG5 and ATG16L1 was significantly increased upon GW4869 treatment in MDA-MB-231 cells (Fig. 3D and HEK293T cells (Fig. 3E). Interestingly, other autophagy genes such as ATG7, ATG12, Beclin1 and LC3II didi not show significant change in expression upon GW4869 treatment, thus highlighting the unique role of ATG5 and ATG16L1 in exosome pathway. Increase in ATG5 protein expression was also observed with different concentrations of GW4869 (Supp. Fig. 3B). GW4869 is an inhibitor of nSmase2 and we observed a negative correlation of SMPD3 gene (nSmase2) with ATG5 and ATG16L1 in both normal and cancer databases, thus validating the reciprocal relation between nSmase 2 and expression of ATG5 and ATG16L1. (Supp. Fig. 3C).

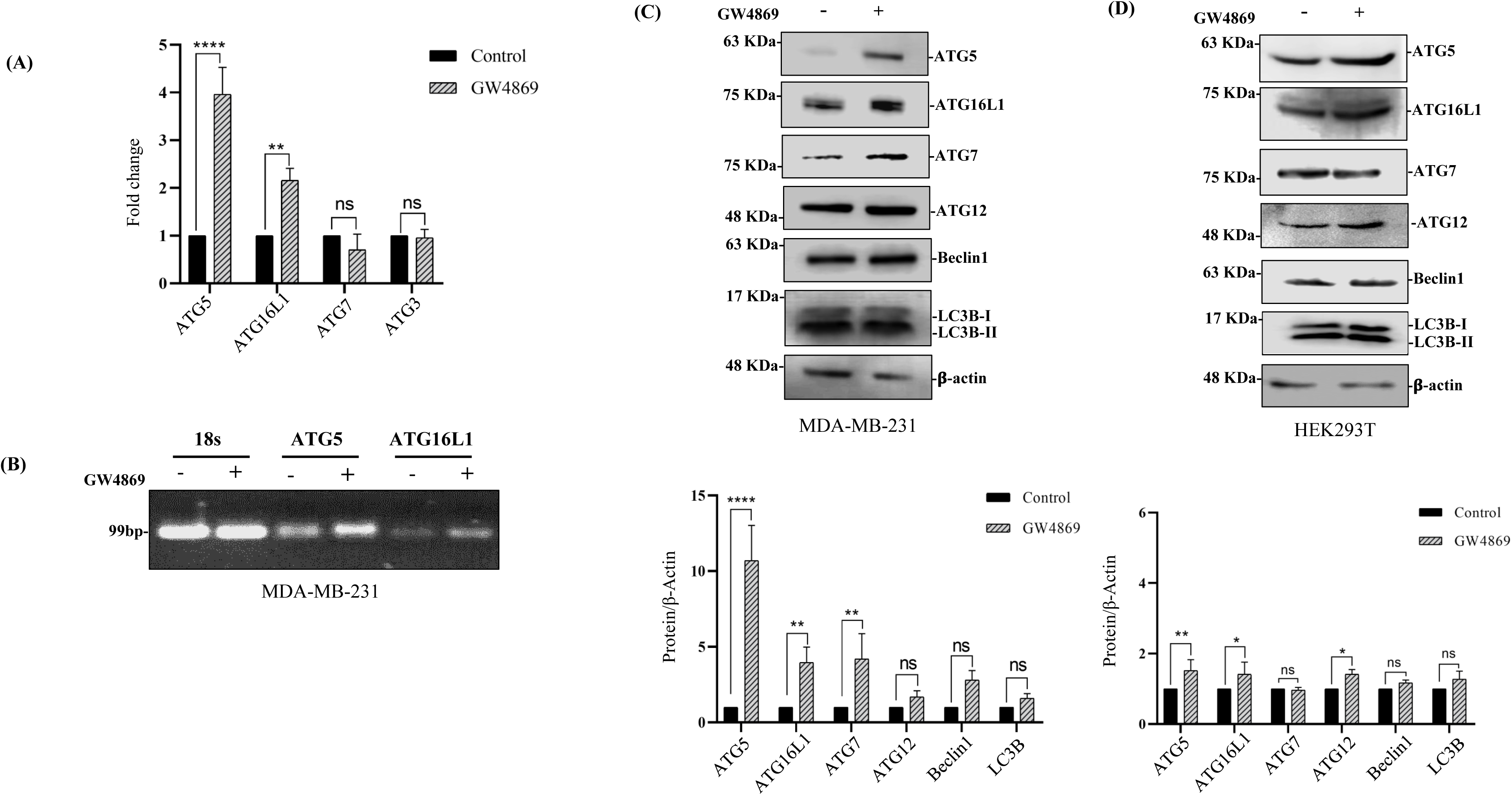

### 4. Combined inhibition of autophagy and exosome biogenesis induces cell death in breast cancer cells

Previous studies have shown that GW4869 potentially inhibits tumor growth in mouse 4T1 breast tumor model system[46]. On contrary, studies have also shown that nSmase2 (SMPD3) inhibits cell proliferation and stops the growth of drug-resistant tumors[47], [48]. So, we performed a cell proliferation assay and found that treatment of MDA-MB-231 cells with 10µM GW4869 for 24 hrs significantly enhances their proliferation (Fig. 4A). In a cell-count assay, we also validated the effect of GW4869 on the MDA-MB-231 proliferation rate. As compared to treatment with CQ and wortmannin, treatment with GW4869 led to increase in the cell number (Supp. Fig. 4A). Following these results, we treated MDA-MB-231 cells with increasing concentration of GW4869 (2.5µM, 5µM, and 10µM) and a combination treatment of 10µM CQ+10µM GW4869 for 24 hr (Fig. 4B). We observed that GW4869 treatment alone gradually increases cell proliferation but combined treatment with 10µM CQ caused significant cell death in MDA-MB-231. Similar results were also obtained when the same concentrations were used for MCF-7 cell line (Supp. Fig 4B). The sharp decline in cell viability of MDA-MB-231 and MCF-7 cells upon combined inhibition of autophagy flux and exosome biogenesis highlights the crosstalk between autophagy and exosome pathways and their importance for cancer cell survival. Survival analysis for the SMPD3 gene shows that high expression of SMPD3 increases overall survival in cervical cancer patients, supporting a tumor suppressive role (Supp. Fig. 4C). Further, to validate the effect of combined treatment on Alix expression, we analyzed the Alix expression in MDA-MB-231 and HEK293T cells. Figures 4C and 4D show that the CQ treatment decreases Alix expression. However, a combined treatment of CQ and GW4869 restored the Alix expression. This result indicates that inhibition of autophagy flux by CQ enhances the encapsulation and release of Alix in exosomes, but blocking exosome biogenesis by GW4869 stops the encapsulation of Alix in exosomes and thus rescues the levels of Alix. This conclusion was also correlated with the TCGA sample databases where PDCD6IP (Alix) showed a progressively decreased expression in luminal, HER2 (+), and TNBC, respectively which might be due to enhanced autophagy perturbation in these tumor types (Supp. Fig. 4D). This result highlights the dynamic crosstalk between autophagy and exosome pathways and potential of restricting cancer cell growth by combined inhibition of the two pathways simultaneously.

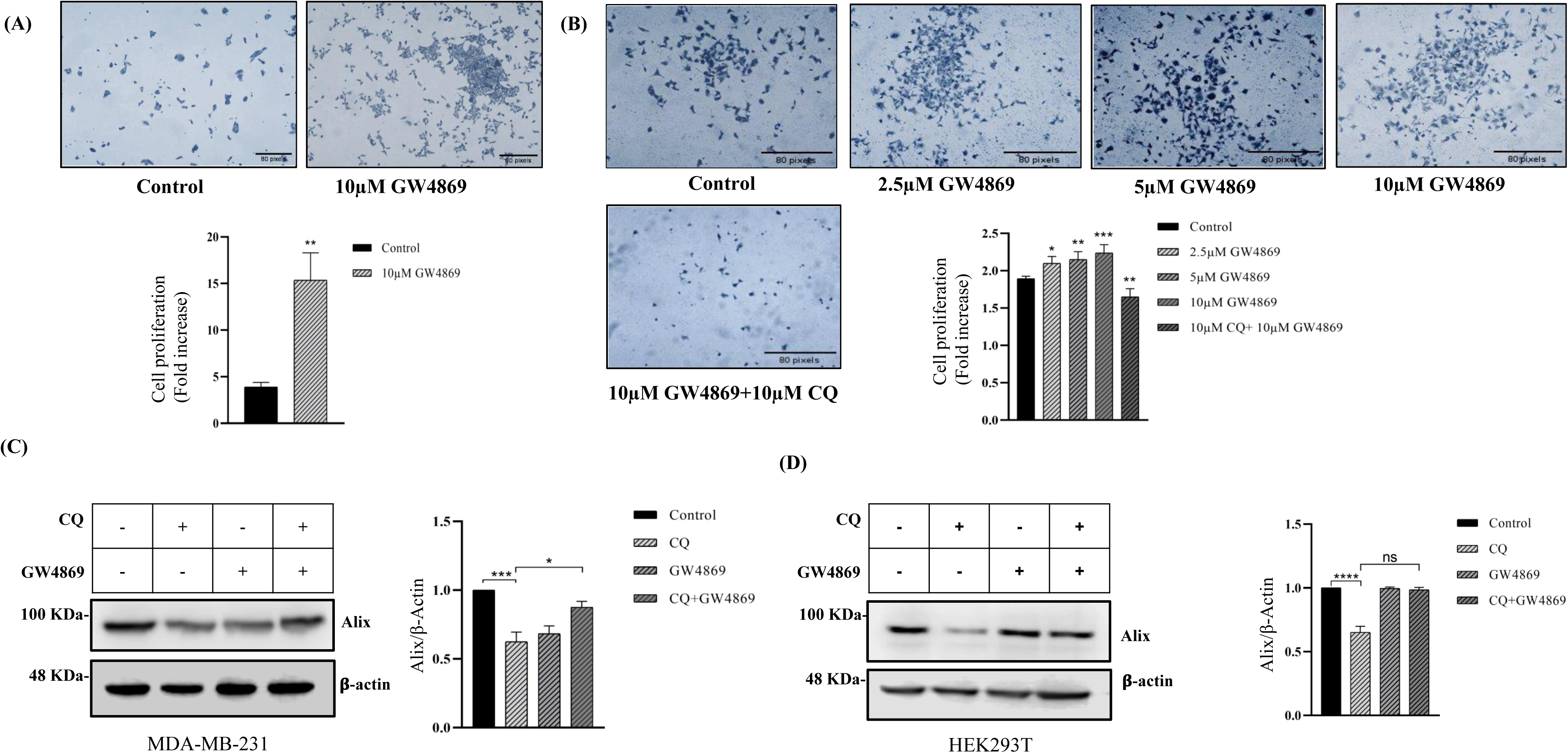

### 5. Inhibition of exosome biogenesis by GW4869 alters the expression of ATG8 family genes

Previous studies have identified an autophagy-mediated unconventional protein secretion responsible for release of molecules not secreted through conventional protein secretion[49]. The unconventional secretion by autophagy may partially be mediated through exosome pathway. It is unclear whether ATG8s particularly GABARAP secretion through exosomes was either a by-product of cellular trafficking or a specific event [50], [51]. Earlier, we assessed the expression of the MAP1LC3 family in GW4869-treated MDA-MB-231 cells, which did not show any significant changes in both MAP1LC3A and MAP1LC3B in GW4869-treated HEK293T cells (Fig. 5C). So, we further checked the expression of another class of ATG8 genes, i.e. GABARAP family. We initially assessed the GABARAPs expression profile in the exoRBase database. We found that GABARAPL2 was highly secreted in different cancer exosomes among other members (Supp. Fig. 5A). Furthermore, the expression of GABARAPL2 showed a significantly higher expression in BRCA than the benign tumors (Supp. Fig. 5B, 5C). Previous studies on EV proteomics have described the association of GABARAP proteins with EV proxitome [51], [52]. A recent study has also demonstrated that lipidated GABARAPs interact with the ESCRT machinery, specifically with the Alix [53]. Since GABARAPL2 showed significantly high expression in BRCA tumors, we performed a real-time PCR for GABARAP genes to check their expression after GW4869 treatment. The results showed that GABARAPL2 expression was significantly decreased after treatment with 10µM GW4869 in MDA-MB-231 (Fig. 5A) as well as HEK293T cells (Fig. 5B). However, no significant changes were observed in both GABARAP and GABARAPL1 genes. These changes were retained in their protein levels where GABARAPL2 expression was significantly decreased in GW4869 treated HEK293T cells (Fig. 5D). This result highlights a potential involvement of GABARAPL2 in exosome pathway

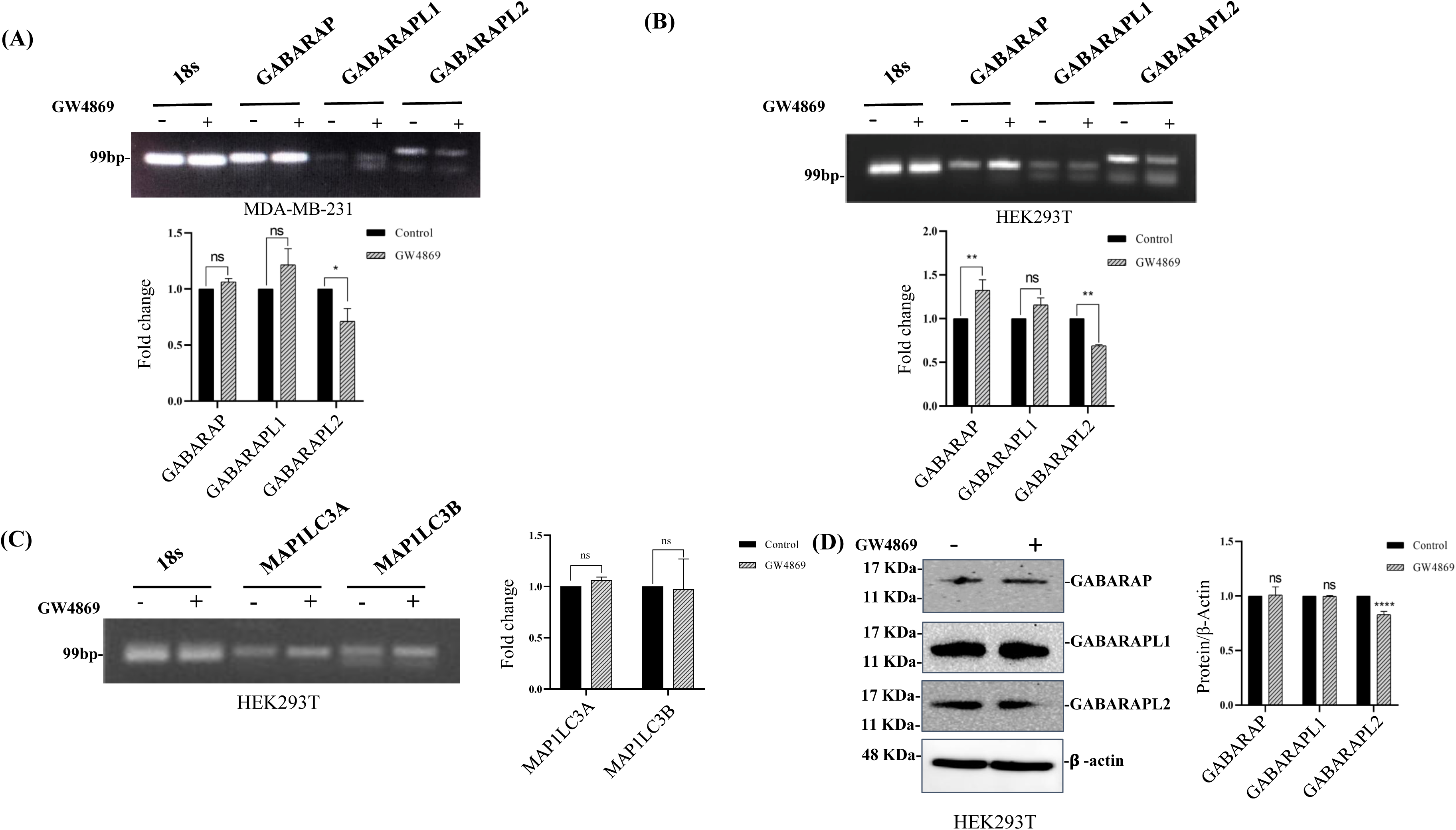

### 6. GABARAPL2 knockdown reversed the CQ-mediated decrease in Alix expression

Previous studies have revealed that ATG12-ATG3 and Alix interact together to regulate endosomal machinery for exosome biogenesis[54]. To check whether GABARAPs are associated with the exosome pathway, we knocked down GABARAP and GABARAPL1 and GABARAPL2 and checked the effect on Alix expression. Interestingly, we observed that the knockdown of GABARAPL2 rescued the CQ-mediated decrease in Alix expression at mRNA and protein levels (Fig. 6A, 6B). However, there was no measurable change in the Alix protein levels after knockdown of GABARAP and GABARAPL1 and CQ treatment (Supp. Fig. 6A). Further, overexpression of GABARAP, GABARAPL1 and GABARAPL2 did not significantly affected the levels of Alix levels (Supp. Fig. 6C). We could conclude that the decrease in Alix expression by CQ is dependent on GABARAPL2. To ascertain the involvement of GABARAPL2 in Alix trafficking in ILVs and subsequent release in exosomes, we analyzed the interaction between Alix and GABARAPL2 using ClusPro, and HDock and visualized by Pymol[55], [56], [57]. To assess this, the following PDB IDs were used to get crystallographic structures: 2OEV-Alix, 4CO7-GABARAPL2. We observed that the HDock Docking score for the Alix-GABARAPL2 complex was -246.29, and the ClusPro docking score was -641.4. Also, Alix residues ranging from 232 to 290 show a strong relation with GABARAPL2 residues range 56 to 116 (Fig. 6C). Further analysis revealed that GABARAPL2 potentially interacts with the BRO domain of Alix. To confirm this, we performed immunoprecipitation of Alix with YFP labeled GABARAPL2, confirming the intracellular interaction of Alix with GABARAPL2 (Supp. Fig. 6A). In conclusion, it is suggested that GABARAPL2 interacts with the Alix to facilitate its release in the exosomes.

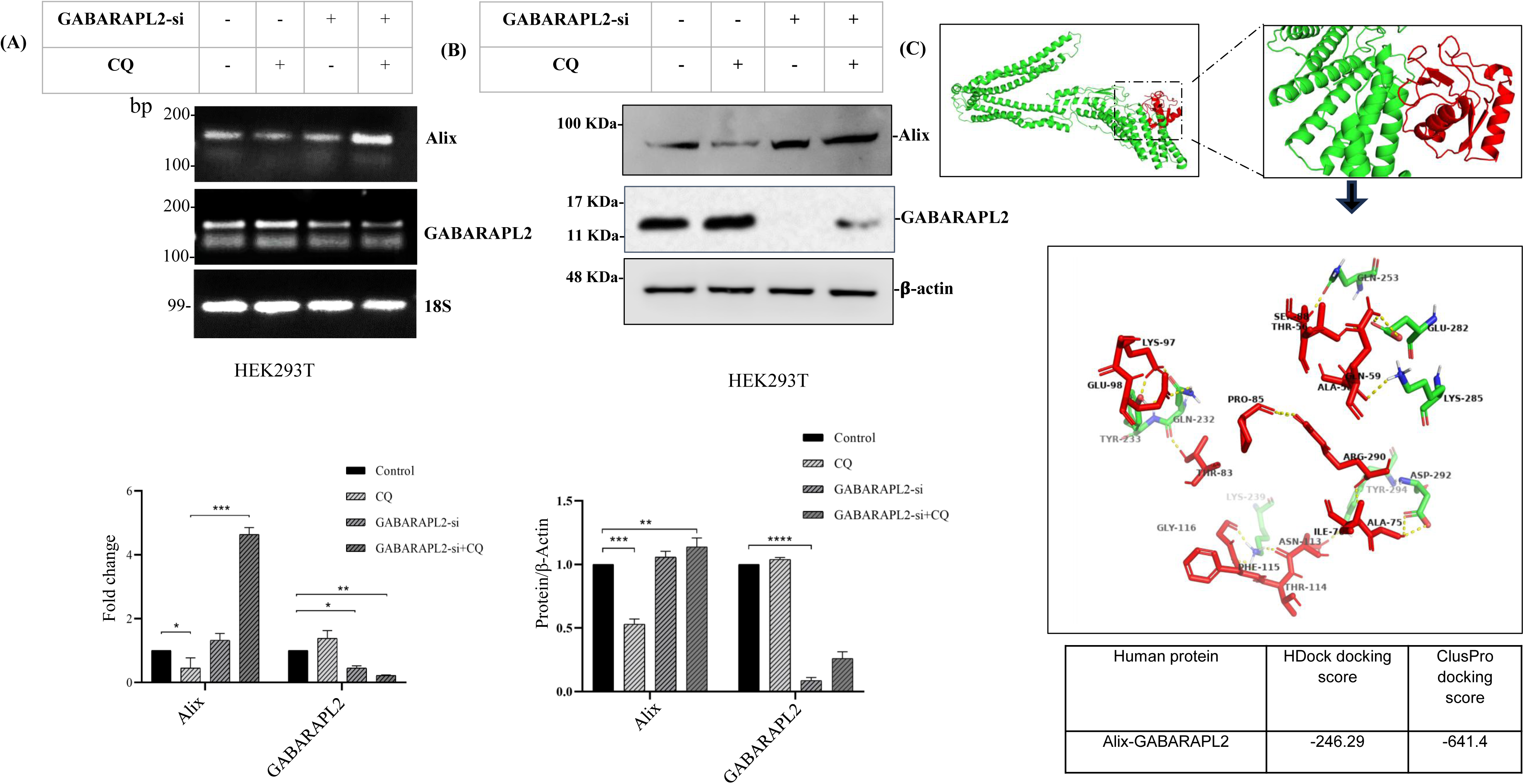

### 7. GABARAPL2 knockdown increases colocalization of Alix and LC3B

We performed confocal imaging to understand the role of GABARAPL2 in modulating Alix puncta. We knocked down GABARAPL2 in MDA-MB-231 cells and then expressed Alix-mNeonGreen and mCherry-LC3B in control, and GABARAPL2 knocked down MDA-MB-231 cells.

After 24 hrs of transfection, cells were treated with 20µM CQ for 24 hours. We observed where average numbers of colocalized puncta were 140 in control, decreased after CQ treatment to 108, and later rescued in combined GABARAPL2 knockdown and CQ to 227 (Fig. 7A and 7B). To understand whether the effect of CQ was specific for Alix localization we checked the puncta of CD63 as well and did not observe any significant change whereas Alix puncta were significantly altered by CQ treatment and rescued by GABARAPL2 knockdown (Supp Fig 7A and 7B). This result highlights the crucial role of GABARAPL2 in directing Alix to exosomes and subsequent release. In the absence of GABARAPL2, Alix remains associated with amphisomes, highlighted by colocalization of Alix and LC3 puncta. Thus, GABARAPL2 facilitates the incorporation of Alix in MVBs and its release in exosomes therefore we conclude that Alix and GABARAPL2 coordinate the crosstalk between autophagy and exosome pathways and these molecules can be targeted for future studies and therapeutic applications.

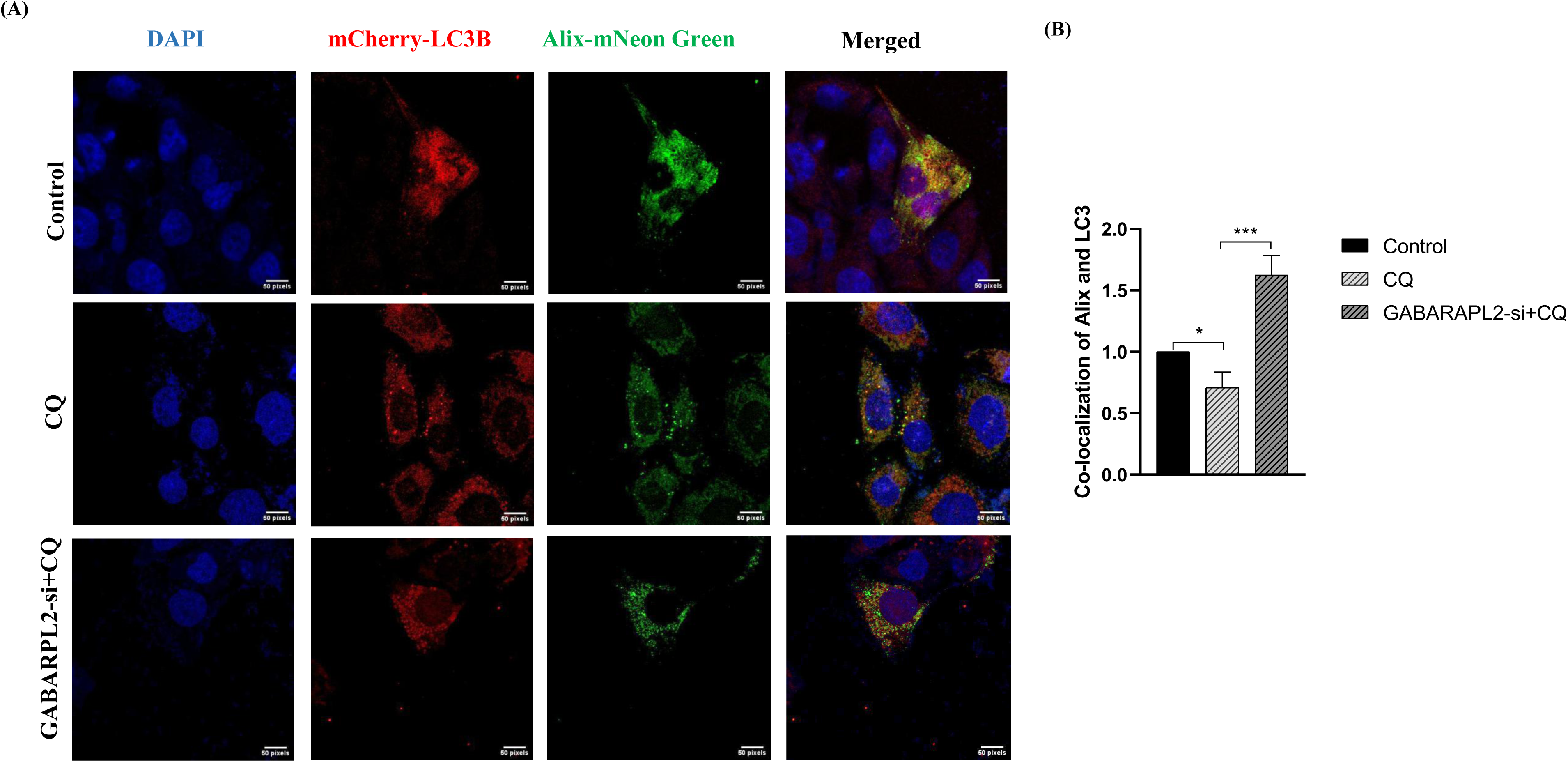

## Discussion

In breast cancer cells, autophagy and exosome biogenesis pathways play an essential role in cellular homeostasis to tolerate hostile conditions in the tumor microenvironment and increase overall survival[58]. In the present study, we wanted to analyze the crosstalk between autophagy and exosome pathways. Unbiased quantitative expression profiling experiments have been thoroughly studied to characterize exosomes after CQ-induced variations. Our first objective was to find out the association between both pathways. For this, we analyzed the gene sets of autophagy and exosome pathways through HADb, KEGG, UALCAN, GEPIA, Correlation-analyzer, exoRBase and Onco-DB[59], [60], [61], [62], [63]. Previous literature has observed that impairment in lysosomal integrity alters the secretome of breast cancer cells[64]. Similar to previous studies, our results have also shown that the exosomes from CQ-treated MDA-MB-231 and MCF-7 cells increase the size and diameter[65]. We characterized the exosomes and observed that high levels of CD63, TSG101, Alix, Flotillin, and β-actin were present in the exosomes from CQ-treated cells. We extended our study and checked CD63, Alix, Rab11, and Rab27a expression in the CQ-treated MDA-MB-231 and HEK293 cells. Although both CD63 and Alix decreased in CQ-treated cells, increased expression of Rab11 and Rab27a led us to know that more MVBs are secreting out rather than fusing with the autophagosome. We further analyzed the same effect on proteins, and the result showed that both Flotillin and Alix were exported out in the exosomes.

We further investigated the fact that GW4869 highly affected autophagy-related genes. Related studies have also shown that the knockdown of SMPD3 enhances p62 and LC3 gene expression[66]. However, our study also determined a novel result that SMPD3 inhibition increases ATG5 and ATG16L1 expression in both MDA-MB-231 and HEK293T cells, but LC3 expression does not show many alterations. Moreover, the close association of ATG5 and ATG16L1 with SMPD3 might play a key role in escaping cancer cells to different chemotherapies.

Previous literature has discovered an increased level of ATG8 orthologs in the cell secretome[36], [51]. A recent study has also demonstrated the involvement of GABARAPL1 in the EV cargo loading[67]. In our study, we observed that SMPD3 inhibition was directly related to down-expression of GABARAPL2. Surprisingly, Alix protein, which was reduced in CQ treatment, got rescued when we knocked down GABARAPL2 and retained accumulated even in the presence of CQ. This indicates that GABARAPL2 has been directly involved in the secretion of Alix through exosomes. Apart from that, we also observed that where Alix was secreting out in the presence of CQ, it was reversed once we used a combined dose of CQ and GW4869. Furthermore, the colocalization study of LC3B and Alix has shown that the colocalized puncta were decreased in CQ treatment, which was later rescued once the knockdown of GABARAPL2 was combined with CQ.

CQ has been used widely, combined with other drugs and therapies, in cancer treatment. Although CQ is recognized as a late-stage autophagy inhibitor, it also exhibits autophagy-independent effects[68]. Our study emphasizes a new mechanism of treatment in which a combined dose of CQ and GW4869, along with ongoing radiotherapy and chemotherapy, might strongly suppress metastatic properties and intracellular signaling in breast cancer cells. In the field of exosome therapy, there are multiple challenges like low targeting, high heterogeneity, short half-life, and very poor efficiency that limit the use of exosomes in therapeutic attributes[69]. Despite some potential limitations, targeting exosomes and autophagy and the use of exosomes in diagnostic and therapeutic applications can be assessed in upcoming clinical studies[70].

## Conclusion

From this study, we unravel the intricate relationship between autophagy and exosome pathways in the survivorship of breast cancer cells. On one side, where treatment with CQ results in enhancement in exosome size, another side shows the treatment with GW4869 predominantly affects ATG5 and ATG16L1 genes. Moreover, the combined inhibition of autophagy and exosome pathways with CQ and GW4869, respectively, causes an increase in Alix expression and breast cancer cell death in MDA-MB-231 and MCF-7 cells. However, we also observed an increase in the exportation of Alix and Flotillin from MDA-MB-231 cells to its secretome after treatment with CQ. Expression of Rab11 and Rab27a genes, which are majorly associated with fusion and export of MVBs and autophagosomes, also increases with CQ treatment. This concludes that the later-stage autophagy inhibition affects both ESCRT-dependent and ESCRT-independent exosome biogenesis. Our significant findings through this study were finding the link between GABARAPL2 and Alix proteins. The knockdown of GABARAPL2 increases the retention of Alix inside the cell, but reciprocity by overexpression of GABARAPL2 doesn’t affect Alix’s expression. Nevertheless, western blotting and confocal imaging have validated that a combined dose of knockdown of GABARAPL2 with CQ significantly rescues the Alix.

## Material and methods

### A. Cell culture

Breast cancer cell lines MDA-MB-231 and MCF-7 cells and human embryonic kidney cells (HEK293T) were derived from NCCS (National Center for Cell Sciences), Pune, India. To maintain stabilized cell lines, these cells were grown in DMEM (Dulbecco’s modified Eagle Medium, Himedia), supplemented with 10% FBS (Himedia) and 1% antibiotic (Penicillin and Streptomycin) in 5% CO_2_ at 37°C. These cells were maintained and passaged thrice a week. All the cell experiments were performed at 70% confluency.

### B. Inhibitor Treatment

Chloroquine diphosphate salt was supplied by Acros Organics, Thermofisher (A0423470). A stock solution of 10mM was made in PBS to make further diluted working concentrations. Wortmannin (19545-26-7) and GW4869 (6823-69-4) were purchased from Sigma. Since both Wortmannin and GW4869 are weekly soluble in water, their stock solution was formed in DMSO (TC185, Himedia). For inhibitor studies, CQ was used in incomplete media whereas wortmannin and GW4869 were used in complete media.

### C. Cell proliferation assay

Briefly, 2x 10^3^ cells were seeded in triplicates in each well of a 96-well plate with 100µL DMEM complete media. Cells were allowed to adhere properly, and after reaching 70% confluency, we treated cells alone or with a combination of the above-listed compounds for 24 hr. The inhibitor GW4869 was added in 2.5µM, 5µM, and 10µM concentrations, while CQ was added in 10µM final concentrations. In the second group, we mixed 10µM CQ with the 10µM GW4869 to evaluate relative cell survival upon inhibiting late-stage autophagy and ESCRT independent pathway, respectively. After 24 hr, the cells were stained with 200 µL crystal violet stain. The absorbance of the captured stain was analyzed by OD at 592nm.

### D. Exosome Isolation

#### a) Exosome Isolation by Ultracentrifugation

Ultracentrifugation is the most common method for isolating extracellular vesicles for good yield with high efficiency and minimum cost. Cultured MDA-MB-231 cells seeded in a 100mm^2^ culture dish at an initial density of 2 × 10^6^ cells. Once cells reach ∼70% confluency, they are treated with 10µM CQ in serum-free DMEM media. After 24 hr, an additional 10µM CQ was added to the dishes. For control, cells were grown in DMEM only. After 48 hr, cell culture media (CCM) of grown cells were isolated and immediately subjected to centrifugation at 12,000 x g, 4 °C for 45 minutes to remove cell debris. The supernatant was collected in fresh UC tubes after filtration with 0.22µ filters (Himedia, SF172-50NO) to remove microvesicles and filled with sterile PBS. Ultracentrifugation was done with an MLA-50 (Beckman Coulter) fixed angle rotor at 1,10,000 x g, 4 °C for 2 hours. Obtained pellets were washed with PBS and, ultimately, volume makeup with PBS. A second round of ultracentrifugation was performed at 1,10,000 x g, 4°C for 70 minutes to obtain an exosome pellet. The isolated pellet was dissolved in an adequate amount of PBS for further applications and stored at -80°C.

#### b) Exosome Isolation with ExoQuick kit

CCM collected from MDA-MB-231 cells was centrifuged at 3000 x g for 15 minutes to remove cell debris. The obtained supernatant was collected in a clean tube and mixed with 63µL ExoQuick exosome precipitation solution, inverted the tube to mix well, and incubated this tube straight upright at 4°C. A centrifuge was done at 1500 x g, 4 °C for 30 minutes to get a white exosome pellet. This pellet was washed in PBS and stored at -80 °C for downstream applications.

### E. Exosome characterization

Exosomes isolated from MDA-MB-231 cell culture media were diluted in PBS and processed in a particle-size analyzer (ZEN 5600, DLS) to estimate the size and intensity of exosomes. Nanoparticle tracking analysis (Malvern NanoSight NS300) was also performed for the same sample. The stock was diluted 500 times with PBS to get particles in an equal frame, and precise diameter, concentration, and size distribution were measured.

A pair of exosome samples were fixed with the 2.5% glutaraldehyde, 2% paraformaldehyde, and 0.1M phosphate buffer (pH 7.4) (Fixative solution) at the time of isolation. To get information about morphology and size, exosomes were pelleted down, and a negative stain of phosphotungstic acid was used on the sample. The resultant exosome suspension was applied to the EM grids to promote the distribution of stained exosomes onto the grids. These grids were then washed briefly with PBS to remove excess negative background. The images of purified exosomes were analyzed by TEM (JEM-1400 Flash).

### F. Protein isolation

MDA-MB-231 and HEK293T cells were seeded in 60mm^2^ dishes. After reaching 70% confluency, cells were treated with either chloroquine (20µM), GW4869 (2.5µM, 5µM, 10µM), or a combination of both drugs for 48 hr in only DMEM media. Whole-cell protein was isolated by adding 100µL of NP40 lysis buffer (5M NaCl, 10% NP-40, 1M Tris pH 8.0) with a 1% protease inhibitor cocktail. A colorimetric assay was done by BCA protein assay kit (Thermo Fisher Scientific Inc., #23225, Rockford, IL, USA) to quantify the total protein of MDA-MB-231, HEK293T cells, and exosomes. Assessment of the BCA product was analyzed at 562nm wavelength.

### G. Immunoblot analysis

Briefly, 100µg of protein was loaded into a sodium dodecyl sulfate-polyacrylamide gel electrophoresis (SDS-PAGE). The proteins were transferred onto a PVDF (Immun-Blot, Bio-Rad, # 1620177) membrane at a constant voltage of 65V for 90 minutes. The membranes were blocked for 1 hr using Tris-buffered saline (TBS) with 3% BSA, and then incubated with the primary antibodies against target proteins, such as Alix, Flotillin, ATG5, ATG12, ATG7, ATG16L1, Beclin1, LC3B, GABARAP, GABARAPL1, GABARAPL2, GFP-monoclonal (Invitrogen, GF28R), beta-actin (CST, USA), CD63, and TSG101 (Proteintech) for overnight at 4 °C on the rocker. After washing with TBS-Tween-20 (0.1%), incubation with HRP-conjugated Anti-mouse and Anti-rabbit secondary antibodies (CST, USA) was carried out for 2 hours at room temperature with shaking. Proteins were detected in an enhanced chemiluminescence (ECL) detection kit (Bio-Rad). The obtained data was measured for densitometry graphs using the software ImageJ.

### H. Detection of mRNA levels

MDA-MB-231, MCF-7, and HEK293T cells were seeded at an initial density of 2×10^5^ cells in 35mm^2^ dishes. Once cells reached 70% confluency, they were treated with 20µM CQ, 10µM GW4869, and 200nM Wortmannin. For cDNA synthesis, the extracted RNA was processed with a Verso cDNA synthesis kit (Thermo Fisher, US). For the quantitative real-time PCR, SYBR Green (Powerup SYBR Green, Thermo Fisher, #A25742) was used and detected with the CFX connect detection system (Bio-Rad). All PCR primers used for this work are listed in Table 1.

**Table 1:**
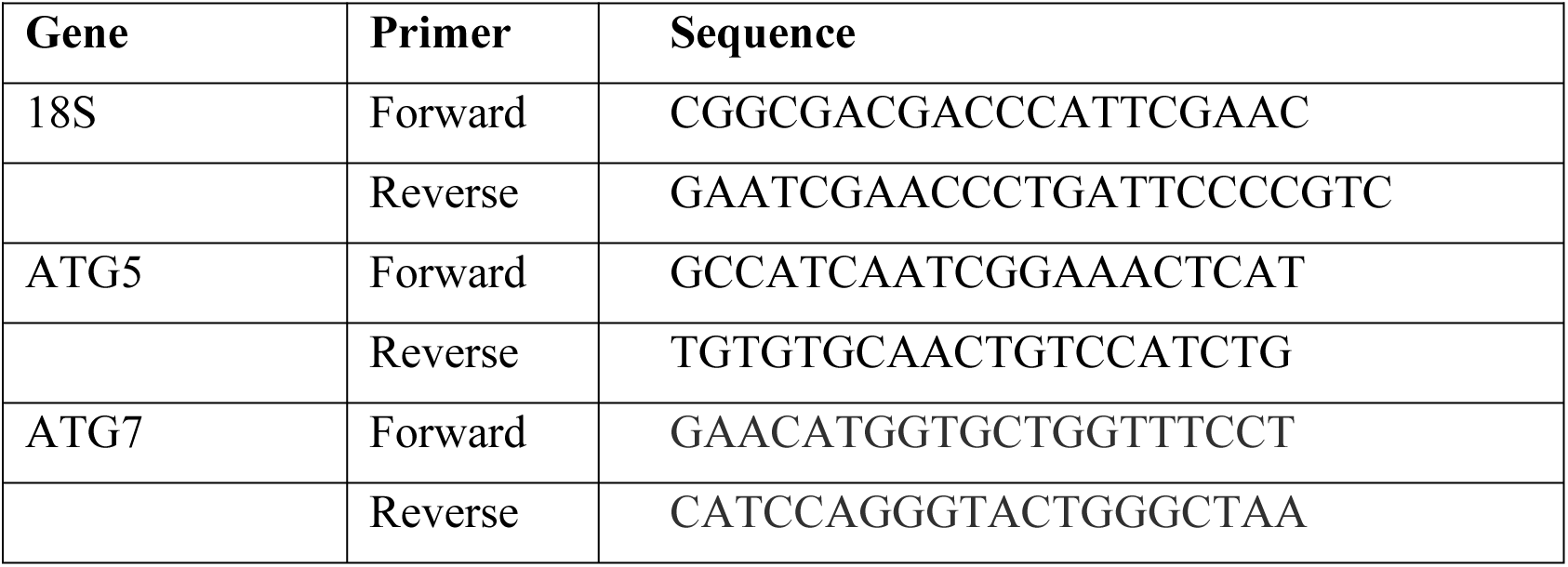
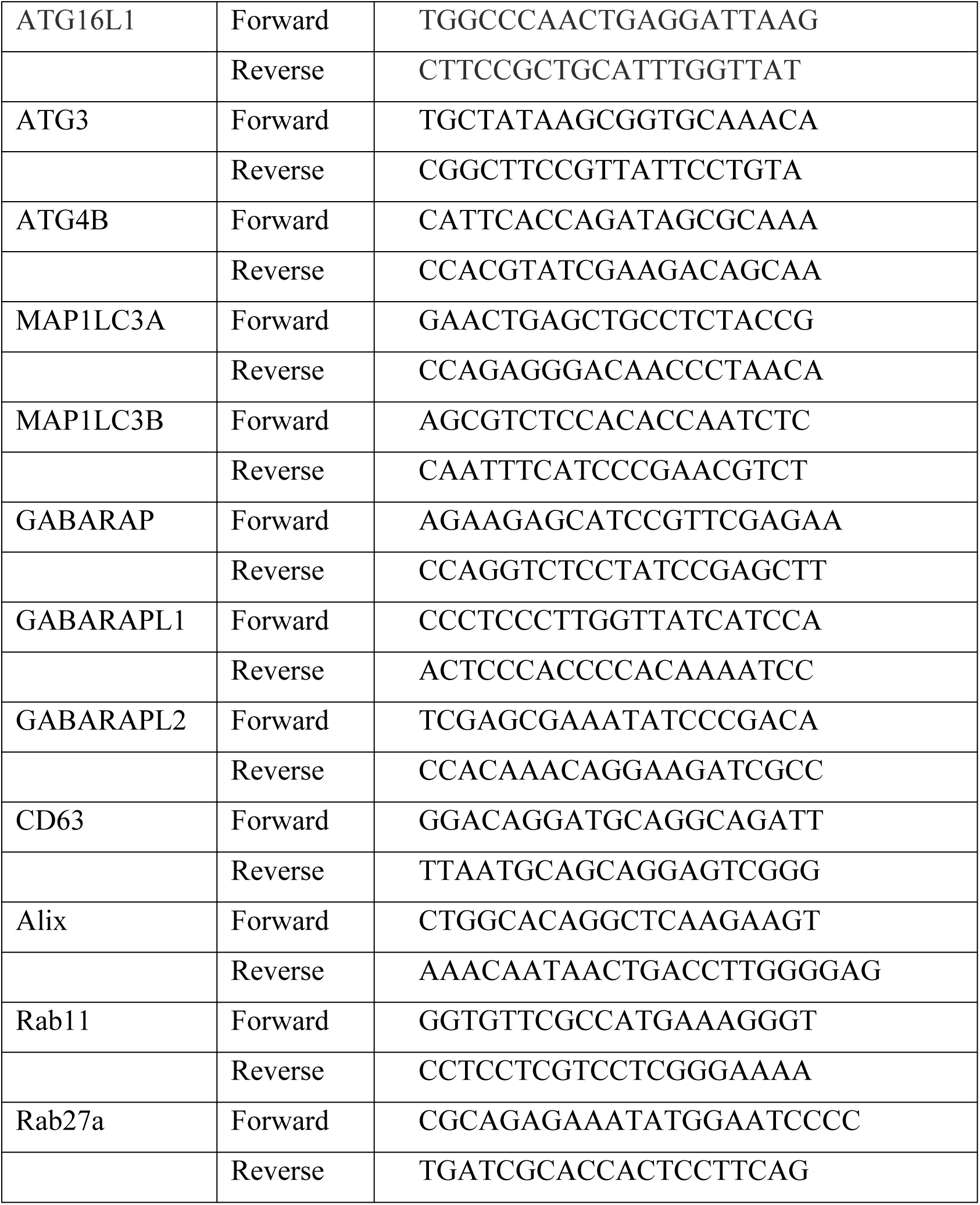
Primer sequences.

### I. Transfection

To knock down the GABARAP isoforms, we used MDA-MB-231 and HEK293T cells and transfected against three different siRNAs, GABARAP (NM_0072782-AS), GABARAPL1 (NM_0314124-AS), and GABRAPL2 (NM_0072857) at 60-70% confluency by using Lipofectamine-3000 (Thermo Fisher Scientific, L3000008) according to the manufacturer’s protocol. Briefly, lipofectamine was used 4µL for a 6-well plate with 20-50nM siRNA. After 24 hours, the media was changed, and the cells were re-transfected with the same siRNA and procedure to get better outcomes. After 24 hours of the second transfection, RNA was isolated and processed into cDNA for further applications. The same protocol was followed for the transfection-based immunoblot studies. For the overexpression studies, control (pcDNA5-EYFP, #47329), GABARAP (pcDNA5/FRT/TO eYFP-GABARAP), GABARAPL1 (pEYFP-GABARAPL1), and GABARAPL2 (pEYFP-hGABARAPL2) plasmids were a kind gift by Dr. Silke Hoffmann Julich, Heinrich Heine University, Germany.

### J. Confocal Fluorscence Analysis

MDA-MB-231 cells were plated in 6 well plates with a seeding density of 2 x 10^4^ cells/well. After reaching 40-50% confluency, knockdown experiments were performed with Lipofectamine 3000. After 24 hours, cells were co-transfected with the mCherry-LC3B and Alix-mNeonGreen plasmids. After 24 hours, CQ treatments were given for another 24 hours. Then, cells were fixed with the 4% paraformaldehyde, washed thrice with PBS, and mounted coverslips with the 70% glycerol. Fluorescence was measured using confocal microscopy (Leica).

Confocal microscopy for the fluorescent-fused proteins was conducted with the Stellaris 8 (Leica) equipped with 63x/1.40 and 100x/1.40 plan achromat objective (oil). For the excitation of DAPI, GFP, mCherry, and mNeonGreen, Argon ion lasers of 405nm, 488nm, 561nm, and 506nm respectively. Smart gain, intensity, and the value of γ were kept optimum throughout the imaging. Images were obtained with LAS-X and later processed and analyzed for colocalization with Fiji[71]. All the images were statistically analyzed using an ANOVA test.

### K. Molecular Docking

We performed protein-protein docking by taking X-ray crystallographic structures for Alix, GABARAP, GABARAPL1, and GABARAPL2. The PDB structures of each GABARAPs (Ligand) were docked with the Alix (Receptor) using Cluspro and HDOCK. The HDOCK is a highly integrated suite that performs on a hybrid algorithm. Cluspro works on a Fast-Fourier Transform (FFT)-based algorithm, enabling it to dock proteins without prior information about the complexes. The final visualization tool used for the complex of docked PDB files was Pymol. This is a molecular visualization software that enables users to carry out virtual screening, binding site prediction, and molecular-docking analysis.

### L. Statistical analysis

All data analyses were done using GraphPad Prism 8.0. We used multiple tests, such as One-way ANOVA, Two-way ANOVA, and t-tests, depending on variables to estimate the difference between groups. A p-value of <0.05 were considered as significant in whole study (*=p<0.05, **=p<0.01, ***=p<0.001, ****=p<0.0001). All the experiments were performed in triplicates.

## Acknowledgment

This work was funded and supported by the Science and Engineering Research Board (SERB)-Power” Grant (SPG/2021/002833), ICMR “Extra mural research grant” (52/27/2020-BIO/BMS) for providing research funds to Dr. Bhawana Bissa.

## Credit Statement

**Naveen Soni:** Conceptualization, writing, Figure preparation, Experimental work**; Megha Chaudhary:** Writing, editing, Experimental work, Proofreading**; Bhawana Bissa:** Conceptualization, writing, supervision

## Conflict of Interest

Authors declare no conflict of interest.

## Abbreviations

ATG: Autophagy-Related
MVBs: Multivesicular Bodies
GABARAP: γ-aminobutyric acid receptor-associated protein
ESCRT: Endosomal Sorting Complex Required for Transport
CQ: Chloroquine

---

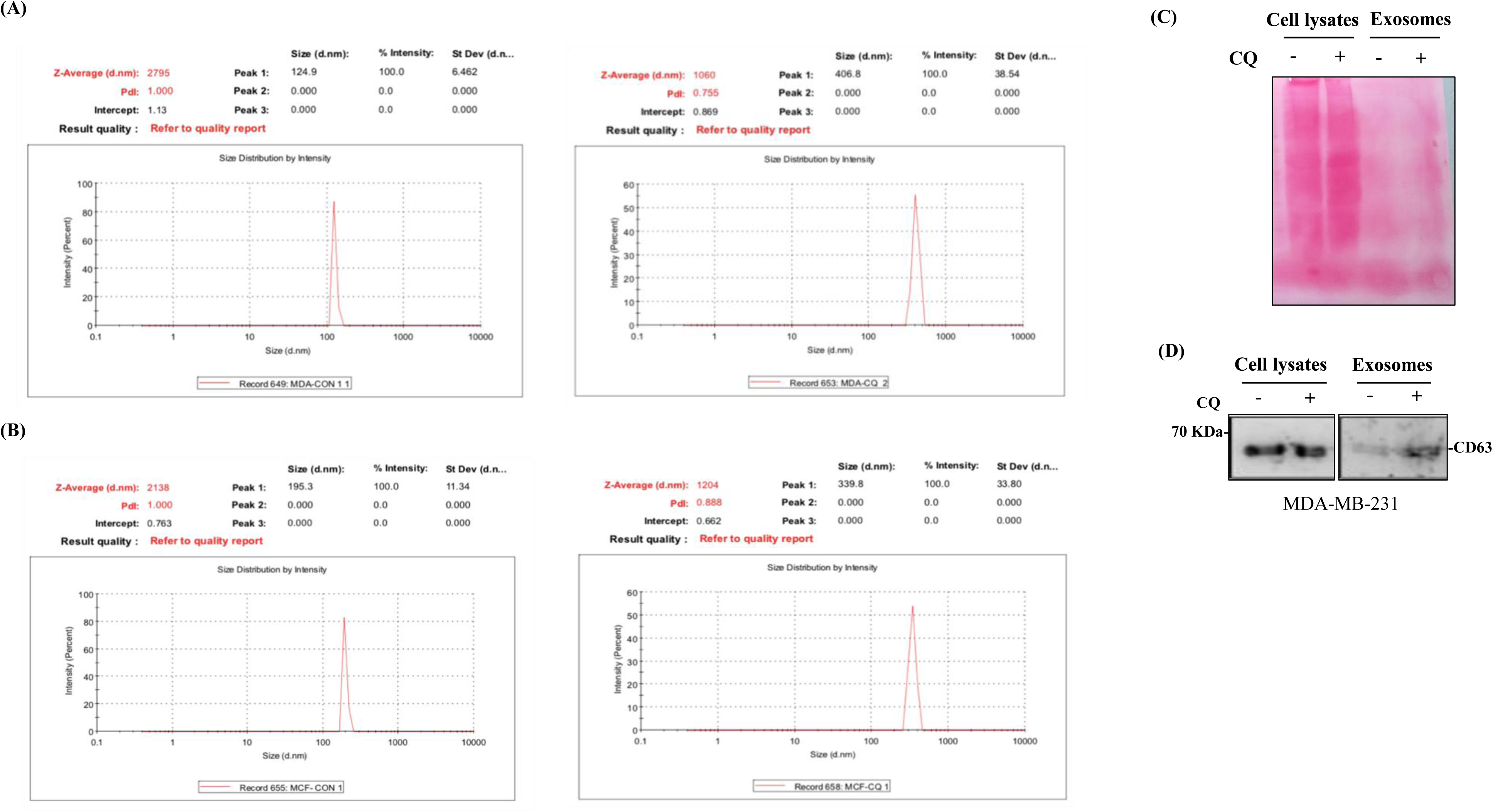

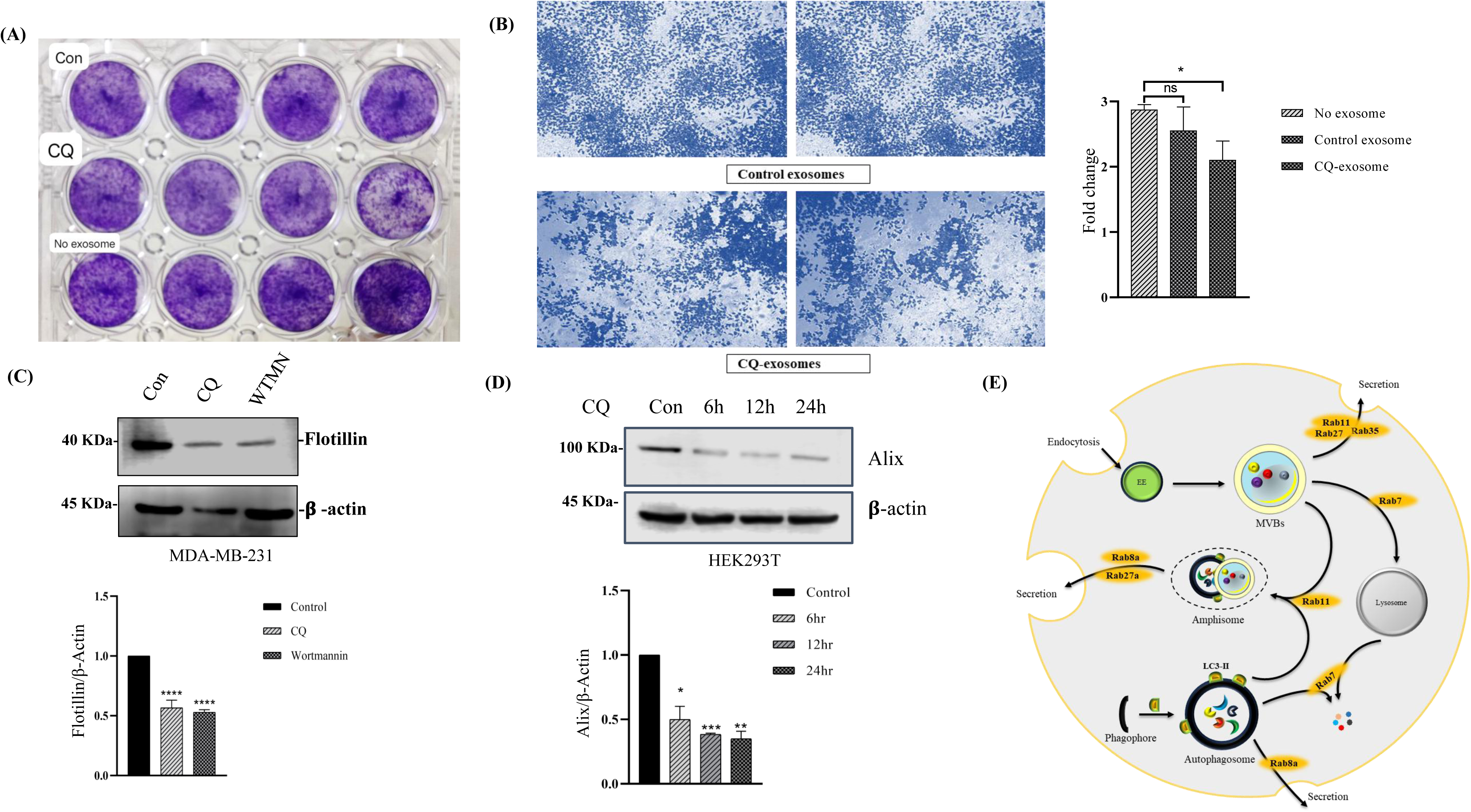

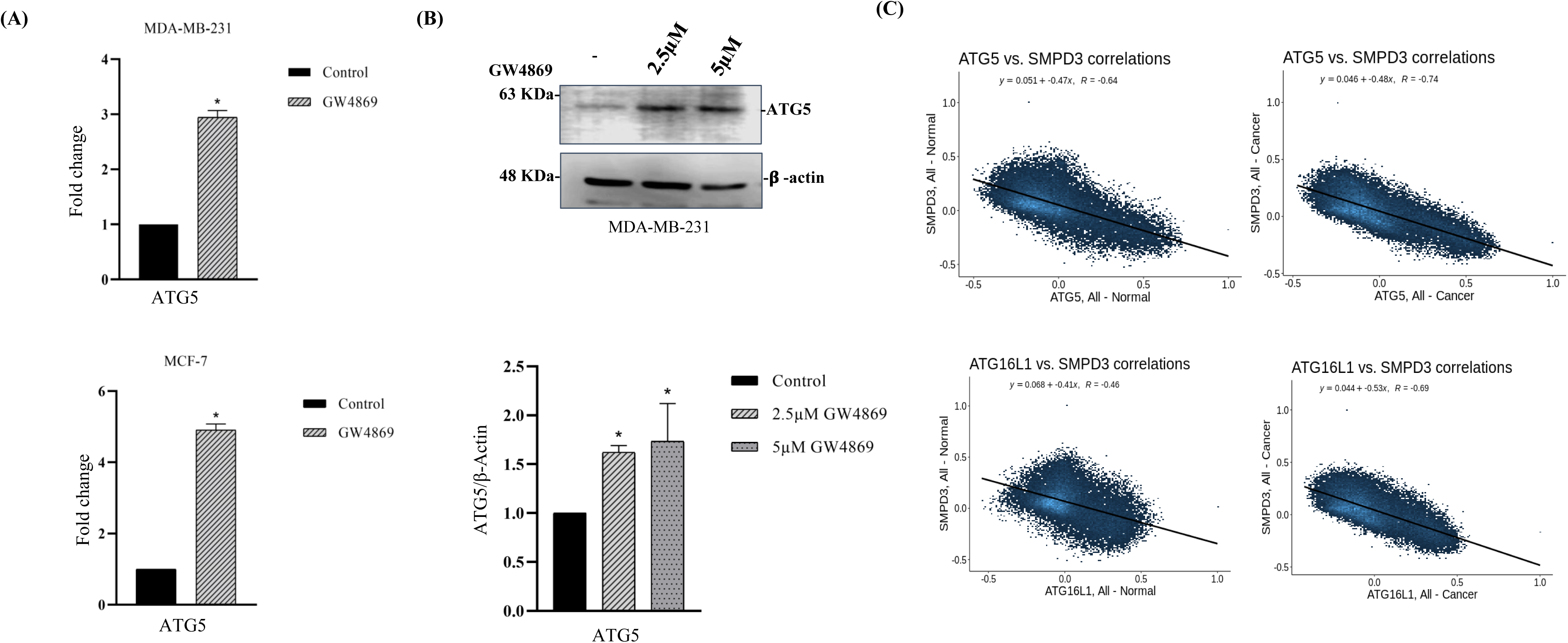

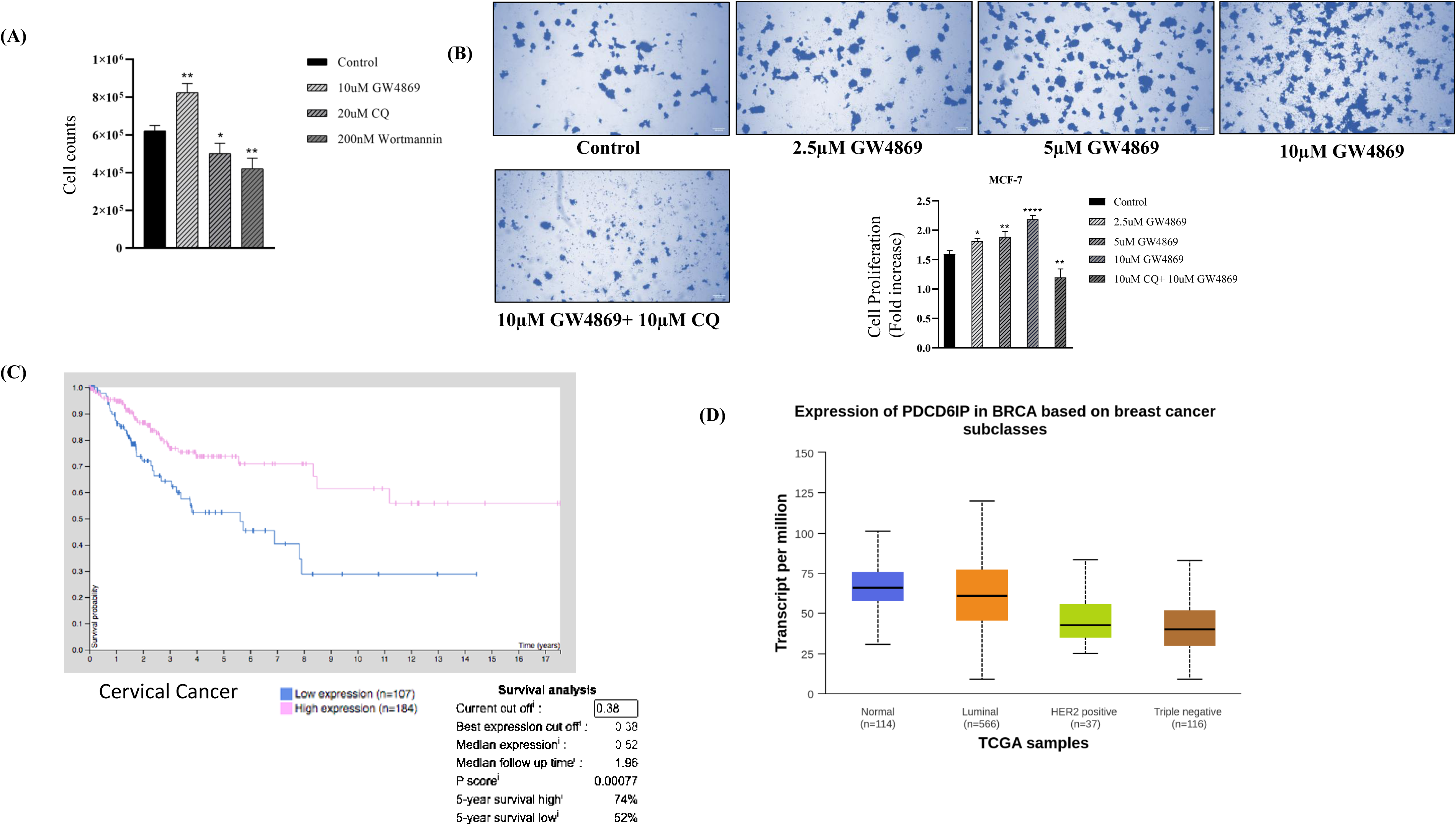

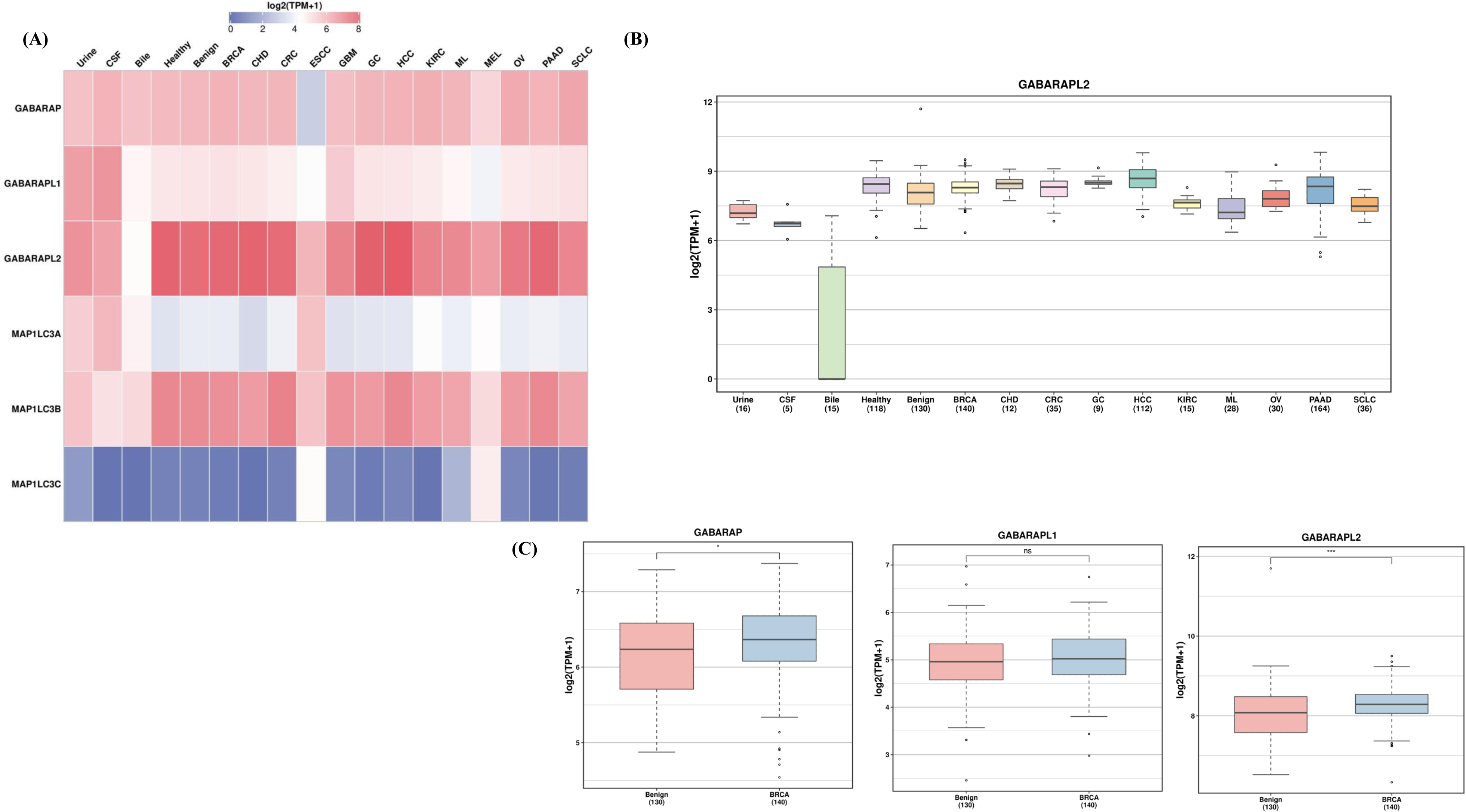

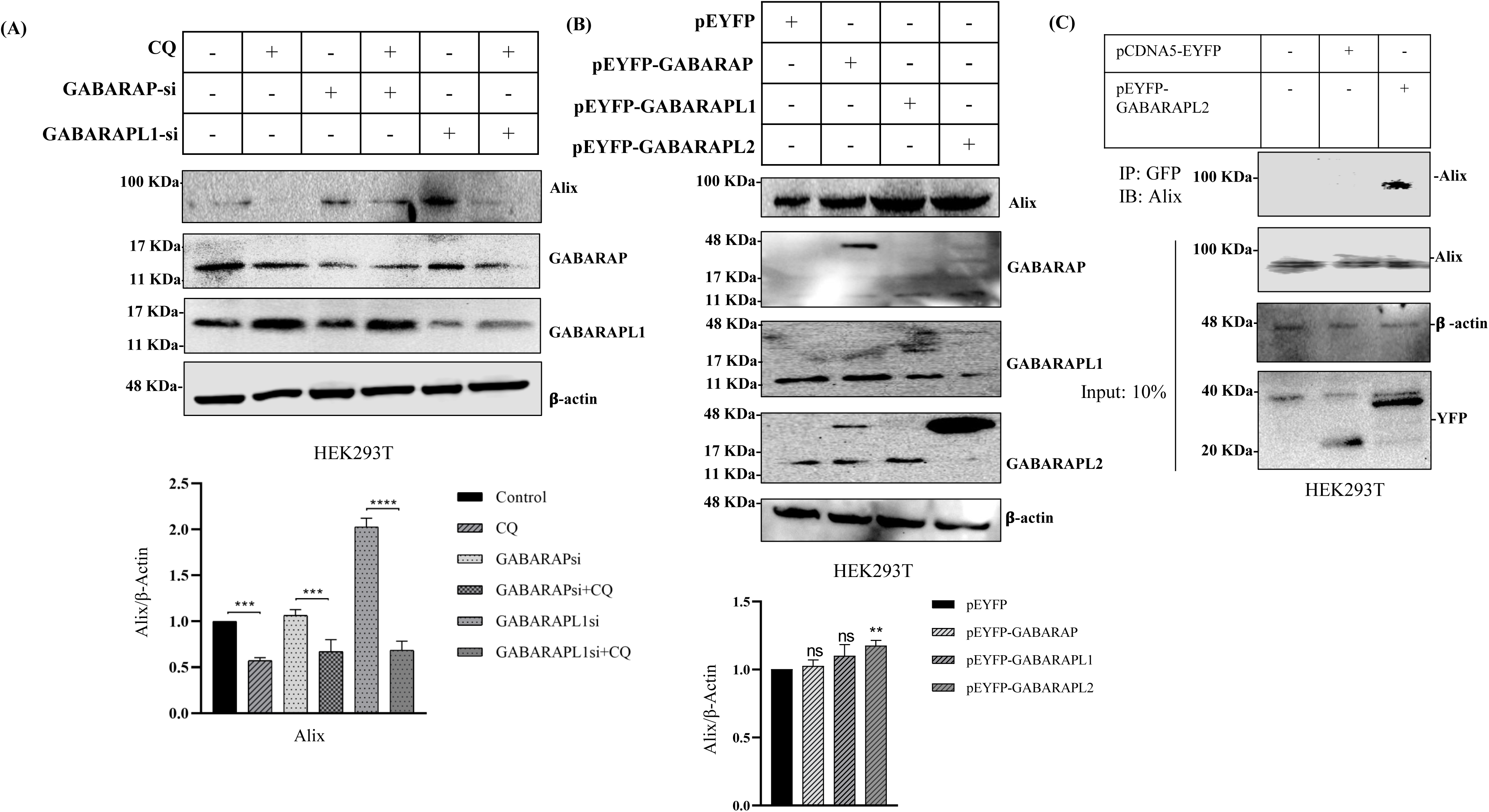

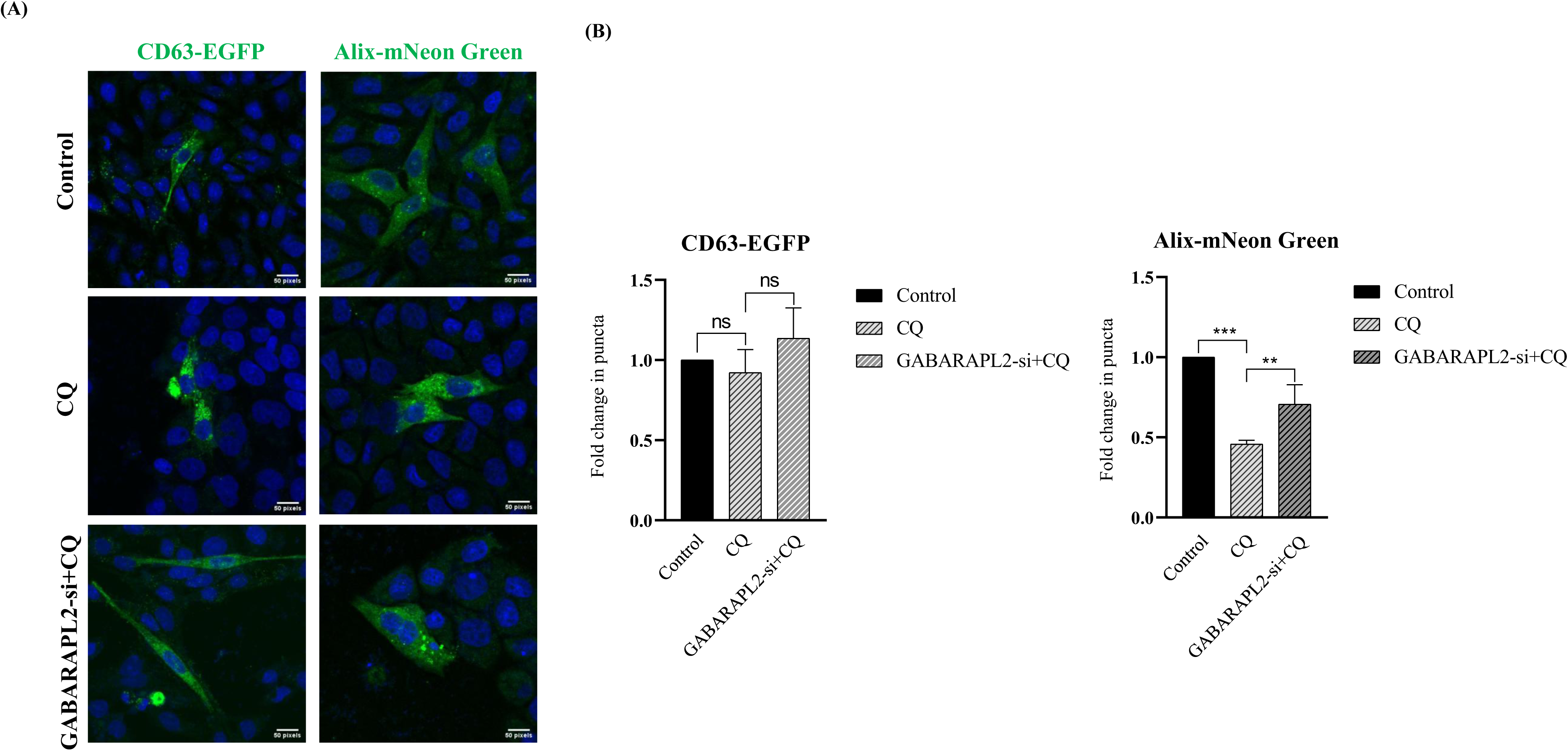

## Notes

### Competing Interest Statement

The authors have declared no competing interest.

## References

[1] D. Bl and C. Bt, “Membrane vesicle release in bacteria, eukaryotes, and archaea: a conserved yet underappreciated aspect of microbial life,” Infection and immunity, vol. 80, no. 6, Jun. 2012, doi: 10.1128/IAI.06014-11.

[2] D. G. Robinson, Y. Ding, and L. Jiang, “Unconventional protein secretion in plants: a critical assessment,” Protoplasma, vol. 253, no. 1, pp. 31–43, Jan. 2016, doi: 10.1007/s00709-015-0887-1.

[3] J. S. Schorey, Y. Cheng, P. P. Singh, and V. L. Smith, “Exosomes and other extracellular vesicles in host-pathogen interactions,” EMBO Rep, vol. 16, no. 1, pp. 24–43, Jan. 2015, doi: 10.15252/embr.201439363.

[4] B. T. Pan and R. M. Johnstone, “Fate of the transferrin receptor during maturation of sheep reticulocytes in vitro: selective externalization of the receptor,” Cell, vol. 33, no. 3, pp. 967–978, Jul. 1983, doi: 10.1016/0092-8674(83)90040-5.

[5] R. M. Johnstone, M. Adam, J. R. Hammond, L. Orr, and C. Turbide, “Vesicle formation during reticulocyte maturation. Association of plasma membrane activities with released vesicles (exosomes).,” Journal of Biological Chemistry, vol. 262, no. 19, pp. 9412–9420, Jul. 1987, doi: 10.1016/S0021-9258(18)48095-7.

[6] G. Raposo and W. Stoorvogel, “Extracellular vesicles: exosomes, microvesicles, and friends,” J Cell Biol, vol. 200, no. 4, pp. 373–383, Feb. 2013, doi: 10.1083/jcb.201211138.

[7] E. Cocucci and J. Meldolesi, “Ectosomes and exosomes: shedding the confusion between extracellular vesicles,” Trends Cell Biol, vol. 25, no. 6, pp. 364–372, Jun. 2015, doi: 10.1016/j.tcb.2015.01.004.

[8] Y. Zhang, Y. Liu, H. Liu, and W. H. Tang, “Exosomes: biogenesis, biologic function and clinical potential,” Cell & Bioscience, vol. 9, no. 1, p. 19, Feb. 2019, doi: 10.1186/s13578-019-0282-2.

[9] N. A. Garcia, I. Ontoria-Oviedo, H. González-King, A. Diez-Juan, and P. Sepúlveda, “Glucose Starvation in Cardiomyocytes Enhances Exosome Secretion and Promotes Angiogenesis in Endothelial Cells,” PLoS One, vol. 10, no. 9, p. e0138849, 2015, doi: 10.1371/journal.pone.0138849.

[10] J. H. Hurley, “ESCRT complexes and the biogenesis of multivesicular bodies,” Curr Opin Cell Biol, vol. 20, no. 1, pp. 4–11, Feb. 2008, doi: 10.1016/j.ceb.2007.12.002.

[11] F. Arslan et al., “Mesenchymal stem cell-derived exosomes increase ATP levels, decrease oxidative stress and activate PI3K/Akt pathway to enhance myocardial viability and prevent adverse remodeling after myocardial ischemia/reperfusion injury,” Stem Cell Res, vol. 10, no. 3, pp. 301–312, May 2013, doi: 10.1016/j.scr.2013.01.002.

[12] W. M. Henne, N. J. Buchkovich, and S. D. Emr, “The ESCRT Pathway,” Developmental Cell, vol. 21, no. 1, pp. 77–91, Jul. 2011, doi: 10.1016/j.devcel.2011.05.015.

[13] J. Kowal, M. Tkach, and C. Théry, “Biogenesis and secretion of exosomes,” Curr Opin Cell Biol, vol. 29, pp. 116–125, Aug. 2014, doi: 10.1016/j.ceb.2014.05.004.

[14] F. Fatima and M. Nawaz, “Stem cell-derived exosomes: roles in stromal remodeling, tumor progression, and cancer immunotherapy,” Chinese Journal of Cancer, vol. 34, no. 3, p. 46, Sep. 2015, doi: 10.1186/s40880-015-0051-5.

[15] M. F. Baietti et al., “Syndecan-syntenin-ALIX regulates the biogenesis of exosomes,” Nat Cell Biol, vol. 14, no. 7, pp. 677–685, Jun. 2012, doi: 10.1038/ncb2502.

[16] J. F. Nabhan, R. Hu, R. S. Oh, S. N. Cohen, and Q. Lu, “Formation and release of arrestin domain-containing protein 1-mediated microvesicles (ARMMs) at plasma membrane by recruitment of TSG101 protein,” Proc Natl Acad Sci U S A, vol. 109, no. 11, pp. 4146–4151, Mar. 2012, doi: 10.1073/pnas.1200448109.

[17] S. Stuffers, C. Sem Wegner, H. Stenmark, and A. Brech, “Multivesicular endosome biogenesis in the absence of ESCRTs,” Traffic, vol. 10, no. 7, pp. 925–937, Jul. 2009, doi: 10.1111/j.1600-0854.2009.00920.x.

[18] K. Trajkovic et al., “Ceramide triggers budding of exosome vesicles into multivesicular endosomes,” Science, vol. 319, no. 5867, pp. 1244–1247, Feb. 2008, doi: 10.1126/science.1153124.

[19] F. M. Goñi and A. Alonso, “Effects of ceramide and other simple sphingolipids on membrane lateral structure,” Biochim Biophys Acta, vol. 1788, no. 1, pp. 169–177, Jan. 2009, doi: 10.1016/j.bbamem.2008.09.002.

[20] G. van Niel et al., “The tetraspanin CD63 regulates ESCRT-independent and - dependent endosomal sorting during melanogenesis,” Dev Cell, vol. 21, no. 4, pp. 708–721, Oct. 2011, doi: 10.1016/j.devcel.2011.08.019.

[21] A. C. Theos et al., “A lumenal domain-dependent pathway for sorting to intralumenal vesicles of multivesicular endosomes involved in organelle morphogenesis,” Dev Cell, vol. 10, no. 3, pp. 343–354, Mar. 2006, doi: 10.1016/j.devcel.2006.01.012.

[22] K.-H. Lim and L. M. Staudt, “Toll-like receptor signaling,” Cold Spring Harb Perspect Biol, vol. 5, no. 1, p. a011247, Jan. 2013, doi: 10.1101/cshperspect.a011247.

[23] A. Salminen, K. Kaarniranta, and A. Kauppinen, “Beclin 1 interactome controls the crosstalk between apoptosis, autophagy and inflammasome activation: impact on the aging process,” Ageing Res Rev, vol. 12, no. 2, pp. 520–534, Mar. 2013, doi: 10.1016/j.arr.2012.11.004.

[24] X. Li, S. He, and B. Ma, “Autophagy and autophagy-related proteins in cancer,” Molecular Cancer, vol. 19, 2020, doi: 10.1186/s12943-020-1138-4.

[25] C. He and D. J. Klionsky, “Regulation mechanisms and signaling pathways of autophagy,” Annu Rev Genet, vol. 43, pp. 67–93, 2009, doi: 10.1146/annurev-genet-102808-114910.

[26] A. Kihara, Y. Kabeya, Y. Ohsumi, and T. Yoshimori, “Beclin-phosphatidylinositol 3-kinase complex functions at the trans-Golgi network,” EMBO Rep, vol. 2, no. 4, pp. 330–335, Apr. 2001, doi: 10.1093/embo-reports/kve061.

[27] B. Levine, R. Liu, X. Dong, and Q. Zhong, “Beclin orthologs: integrative hubs of cell signaling, membrane trafficking, and physiology,” Trends Cell Biol, vol. 25, no. 9, pp. 533–544, Sep. 2015, doi: 10.1016/j.tcb.2015.05.004.

[28] Y. Xin et al., “Cloning, expression patterns, and chromosome localization of three human and two mouse homologues of GABA(A) receptor-associated protein,” Genomics, vol. 74, no. 3, pp. 408–413, Jun. 2001, doi: 10.1006/geno.2001.6555.

[29] H. He et al., “Post-translational modifications of three members of the human MAP1LC3 family and detection of a novel type of modification for MAP1LC3B,” J Biol Chem, vol. 278, no. 31, pp. 29278–29287, Aug. 2003, doi: 10.1074/jbc.M303800200.

[30] H. Weidberg, E. Shvets, T. Shpilka, F. Shimron, V. Shinder, and Z. Elazar, “LC3 and GATE-16/GABARAP subfamilies are both essential yet act differently in autophagosome biogenesis,” EMBO J, vol. 29, no. 11, pp. 1792–1802, Jun. 2010, doi: 10.1038/emboj.2010.74.

[31] N. Kosaka, H. Iguchi, Y. Yoshioka, F. Takeshita, Y. Matsuki, and T. Ochiya, “Secretory mechanisms and intercellular transfer of microRNAs in living cells,” J Biol Chem, vol. 285, no. 23, pp. 17442–17452, Jun. 2010, doi: 10.1074/jbc.M110.107821.

[32] A. A. Shamseddine, M. V. Airola, and Y. A. Hannun, “Roles and regulation of neutral sphingomyelinase-2 in cellular and pathological processes,” Adv Biol Regul, vol. 57, pp. 24–41, Jan. 2015, doi: 10.1016/j.jbior.2014.10.002.

[33] Y. Y. Li and S. J. Jones, “Drug repositioning for personalized medicine,” Genome Medicine, vol. 4, no. 3, p. 27, Mar. 2012, doi: 10.1186/gm326.

[34] C. Verbaanderd et al., “Repurposing Drugs in Oncology (ReDO)-chloroquine and hydroxychloroquine as anti-cancer agents,” Ecancermedicalscience, vol. 11, p. 781, 2017, doi: 10.3332/ecancer.2017.781.

[35] V. R. Solomon and H. Lee, “Chloroquine and its analogs: a new promise of an old drug for effective and safe cancer therapies,” Eur J Pharmacol, vol. 625, no. 1–3, pp. 220–233, Dec. 2009, doi: 10.1016/j.ejphar.2009.06.063.

[36] J. Xu et al., “Chloroquine treatment induces secretion of autophagy-related proteins and inclusion of Atg8-family proteins in distinct extracellular vesicle populations,” Autophagy, vol. 18, no. 11, pp. 2547–2560, doi: 10.1080/15548627.2022.2039535.

[37] M. Colombo, G. Raposo, and C. Théry, “Biogenesis, secretion, and intercellular interactions of exosomes and other extracellular vesicles,” Annu Rev Cell Dev Biol, vol. 30, pp. 255–289, 2014, doi: 10.1146/annurev-cellbio-101512-122326.

[38] C. Théry et al., “Minimal information for studies of extracellular vesicles 2018 (MISEV2018): a position statement of the International Society for Extracellular Vesicles and update of the MISEV2014 guidelines,” J Extracell Vesicles, vol. 7, no. 1, p. 1535750, 2018, doi: 10.1080/20013078.2018.1535750.

[39] F. G. Ortega et al., “Interfering with endolysosomal trafficking enhances release of bioactive exosomes,” Nanomedicine: Nanotechnology, Biology, and Medicine, vol. 20, Aug. 2019, doi: 10.1016/j.nano.2019.102014.

[40] S. S. Panda, R. K. Sahoo, S. K. Patra, S. Biswal, and B. K. Biswal, “Molecular insights to therapeutic in cancer: role of exosomes in tumor microenvironment, metastatic progression and drug resistance,” Drug Discovery Today, vol. 29, no. 8, p. 104061, Aug. 2024, doi: 10.1016/j.drudis.2024.104061.

[41] Y. Huang et al., “Exosome-mediated remodeling of the tumor microenvironment: From local to distant intercellular communication,” Cancer Lett, vol. 543, p. 215796, Sep. 2022, doi: 10.1016/j.canlet.2022.215796.

[42] M. Ostrowski et al., “Rab27a and Rab27b control different steps of the exosome secretion pathway,” Nat Cell Biol, vol. 12, no. 1, pp. 19–30; sup pp 1–13, Jan. 2010, doi: 10.1038/ncb2000.

[43] M. S. Ostenfeld et al., “Cellular disposal of miR23b by RAB27-dependent exosome release is linked to acquisition of metastatic properties,” Cancer Res, vol. 74, no. 20, pp. 5758–5771, Oct. 2014, doi: 10.1158/0008-5472.CAN-13-3512.

[44] H. Guo et al., “Atg5 Disassociates the V1V0-ATPase to Promote Exosome Production and Tumor Metastasis Independent of Canonical Macroautophagy,” Dev Cell, vol. 43, no. 6, pp. 716–730.e7, Dec. 2017, doi: 10.1016/j.devcel.2017.11.018.

[45] N. Fussi et al., “Exosomal secretion of α-synuclein as protective mechanism after upstream blockage of macroautophagy,” Cell Death Dis, vol. 9, no. 7, p. 757, Jul. 2018, doi: 10.1038/s41419-018-0816-2.

[46] B. A. Pulaski and S. Ostrand-Rosenberg, “Mouse 4T1 breast tumor model,” Curr Protoc Immunol, vol. Chapter 20, p. Unit 20.2, May 2001, doi: 10.1002/0471142735.im2002s39.

[47] C. Tallon et al., “Nipping disease in the bud: nSMase2 inhibitors as therapeutics in extracellular vesicle-mediated diseases,” Drug Discov Today, vol. 26, no. 7, pp. 1656–1668, Jul. 2021, doi: 10.1016/j.drudis.2021.03.025.

[48] B. Ghandour, G. Dbaibo, and N. Darwiche, “The unfolding role of ceramide in coordinating retinoid-based cancer therapy,” Biochem J, vol. 478, no. 19, pp. 3621–3642, Oct. 2021, doi: 10.1042/BCJ20210368.

[49] V. Deretic, S. Jiang, and N. Dupont, “Autophagy intersections with conventional and unconventional secretion in tissue development, remodeling and inflammation,” Trends Cell Biol, vol. 22, no. 8, pp. 397–406, Aug. 2012, doi: 10.1016/j.tcb.2012.04.008.

[50] S. Asano et al., “Phospholipase C-related catalytically inactive protein (PRIP) controls KIF5B-mediated insulin secretion,” Biol Open, vol. 3, no. 6, pp. 463–474, May 2014, doi: 10.1242/bio.20147591.

[51] J. L. Sanwald, G. Poschmann, K. Stühler, C. Behrends, S. Hoffmann, and D. Willbold, “The GABARAP Co-Secretome Identified by APEX2-GABARAP Proximity Labelling of Extracellular Vesicles,” Cells, vol. 9, no. 6, Jun. 2020, doi: 10.3390/cells9061468.

[52] C. Lefebvre, R. Legouis, and E. Culetto, “ESCRT and autophagies: Endosomal functions and beyond,” Semin Cell Dev Biol, vol. 74, pp. 21–28, Feb. 2018, doi: 10.1016/j.semcdb.2017.08.014.

[53] M. Ogura et al., “Microautophagy regulated by STK38 and GABARAPs is essential to repair lysosomes and prevent aging,” EMBO Rep, vol. 24, no. 12, p. e57300, Nov. 2023, doi: 10.15252/embr.202357300.

[54] M. L, M. R, and D. J, “ATG12-ATG3 interacts with Alix to promote basal autophagic flux and late endosome function,” Nature cell biology, vol. 17, no. 3, Mar. 2015, doi: 10.1038/ncb3112.

[55] Y. Yan, D. Zhang, P. Zhou, B. Li, and S.-Y. Huang, “HDOCK: a web server for protein-protein and protein-DNA/RNA docking based on a hybrid strategy,” Nucleic Acids Res, vol. 45, no. W1, pp. W365–W373, Jul. 2017, doi: 10.1093/nar/gkx407.

[56] R. E. Rigsby and A. B. Parker, “Using the PyMOL application to reinforce visual understanding of protein structure,” Biochem Mol Biol Educ, vol. 44, no. 5, pp. 433– 437, Sep. 2016, doi: 10.1002/bmb.20966.

[57] D. Kozakov et al., “The ClusPro web server for protein-protein docking,” Nat Protoc, vol. 12, no. 2, pp. 255–278, Feb. 2017, doi: 10.1038/nprot.2016.169.

[58] M. Colletti, D. Ceglie, A. Di Giannatale, and F. Nazio, “Autophagy and Exosomes Relationship in Cancer: Friends or Foes?,” Front Cell Dev Biol, vol. 8, p. 614178, 2020, doi: 10.3389/fcell.2020.614178.

[59] H. Lai et al., “exoRBase 2.0: an atlas of mRNA, lncRNA and circRNA in extracellular vesicles from human biofluids,” Nucleic Acids Res, vol. 50, no. D1, pp. D118–D128, Jan. 2022, doi: 10.1093/nar/gkab1085.

[60] D. S. Chandrashekar et al., “UALCAN: A Portal for Facilitating Tumor Subgroup Gene Expression and Survival Analyses,” Neoplasia, vol. 19, no. 8, pp. 649–658, Aug. 2017, doi: 10.1016/j.neo.2017.05.002.

[61] “HAMdb: a database of human autophagy modulators with specific pathway and disease information - PubMed.” Accessed: Aug. 02, 2024. [Online]. Available: https://pubmed.ncbi.nlm.nih.gov/30066211/

[62] Z. Tang, C. Li, B. Kang, G. Gao, C. Li, and Z. Zhang, “GEPIA: a web server for cancer and normal gene expression profiling and interactive analyses,” Nucleic Acids Res, vol. 45, no. Web Server issue, pp. W98–W102, Jul. 2017, doi: 10.1093/nar/gkx247.

[63] H. E. Miller and A. J. R. Bishop, “Correlation AnalyzeR: functional predictions from gene co-expression correlations,” BMC Bioinformatics, vol. 22, no. 1, p. 206, Apr. 2021, doi: 10.1186/s12859-021-04130-7.

[64] A. Latifkar et al., “Loss of Sirtuin 1 Alters the Secretome of Breast Cancer Cells by Impairing Lysosomal Integrity,” Dev Cell, vol. 49, no. 3, pp. 393–408.e7, May 2019, doi: 10.1016/j.devcel.2019.03.011.

[65] C. H. Mitchell and W. Lu, “Exosome release from RPE cells increased by lysosomal compromise,” Investigative Ophthalmology & Visual Science, vol. 63, no. 7, pp. 4617–F0409, Jun. 2022.

[66] M. J. Back et al., “Activation of neutral sphingomyelinase 2 by starvation induces cell-protective autophagy via an increase in Golgi-localized ceramide,” Cell Death Dis, vol. 9, no. 6, p. 670, Jun. 2018, doi: 10.1038/s41419-018-0709-4.

[67] J. E. J. Beaumont et al., “GABARAPL1 is essential in extracellular vesicle cargo loading and metastasis development,” Radiotherapy and Oncology, vol. 190, p. 109968, Jan. 2024, doi: 10.1016/j.radonc.2023.109968.

[68] P. Maycotte, S. Aryal, C. T. Cummings, J. Thorburn, M. J. Morgan, and A. Thorburn, “Chloroquine sensitizes breast cancer cells to chemotherapy independent of autophagy,” Autophagy, vol. 8, no. 2, pp. 200–212, Feb. 2012, doi: 10.4161/auto.8.2.18554.

[69] S. Sadeghi, F. R. Tehrani, S. Tahmasebi, A. Shafiee, and S. M. Hashemi, “Exosome engineering in cell therapy and drug delivery,” Inflammopharmacology, vol. 31, no. 1, pp. 145–169, 2023, doi: 10.1007/s10787-022-01115-7.

[70] T. Li et al., “The Therapeutic Potential and Clinical Significance of Exosomes as Carriers of Drug Delivery System,” Pharmaceutics, vol. 15, no. 1, p. 21, Dec. 2022, doi: 10.3390/pharmaceutics15010021.

[71] J. Schindelin et al., “Fiji: an open-source platform for biological-image analysis,” Nat Methods, vol. 9, no. 7, pp. 676–682, Jun. 2012, doi: 10.1038/nmeth.2019.

